# Selective expansion of motor cortical projections in the evolution of vocal novelty

**DOI:** 10.1101/2024.09.13.612752

**Authors:** Emily C. Isko, Clifford E. Harpole, Xiaoyue Mike Zheng, Huiqing Zhan, Martin B. Davis, Anthony M. Zador, Arkarup Banerjee

**Affiliations:** Cold Spring Harbor Laboratory, Cold Spring Harbor, NY; Cold Spring Harbor Laboratory School for Biological Sciences, Cold Spring Harbor, NY

## Abstract

Deciphering how cortical architecture evolves to drive behavioral innovations is a long-standing challenge in neuroscience and evolutionary biology. Here, we leverage a striking behavioral novelty in the Alston’s singing mouse (*Scotinomys teguina*), compared to the laboratory mouse (*Mus musculus*), to quantitatively test models of motor cortical evolution. We used bulk tracing, serial two-photon tomography, and high-throughput DNA sequencing of over 76,000 barcoded neurons to discover a specific and substantial expansion (*∼*200%) of orofacial motor cortical (OMC) projections to the auditory cortical region (AudR) and the midbrain periaqueductal gray (PAG), both implicated in vocal behaviors. Moreover, analysis of individual OMC neurons’ projection motifs revealed preferential expansion of exclusive projections to AudR. Our results imply that selective expansion of ancestral motor cortical projections can underlie behavioral divergence over short evolutionary timescales, suggesting potential mechanisms for the evolution of enhanced cortical control over vocalizations—a crucial preadaptation for human language.

## Introduction

The origins of diverse behavioral traits have fascinated biologists for centuries. Such behavioral diversity prompts an essential question in neuroscience: How are neural circuits modified over evolutionary time to generate novel behaviors? The role of neocortex, which is a defining feature of mammalian brain evolution, is particularly relevant for behavioral innovations [1–4]. The overall organizational plan of the cortex is strikingly conserved; however, this gross similarity belies the species-specific cellular and circuit modifications that underlie the enormous diversity of behaviors among mammals [5]. Deciphering the rules by which cortical architecture evolves is key to understanding how it drives behavioral innovations.

Multiple models of how brains evolve at different spatiotemporal scales have been proposed [6–14]. Over long timescales, cortical evolution is likely fueled by changes in the absolute or relative sizes of the cortical fields [15] as well as changes in numbers, composition and spatial distribution of cell-types [16, 17]. Substantial evidence exists for each of these models: the primate cortex compared to that of a rodent shows a massive expansion in cortical field sizes [18, 19], increased dendritic arborization [20, 21] and even a novel interneuron cell-type not found in mice and ferrets [22, 23]. Beyond such changes in brain architecture, other influential models predict that modified long-range connectivity among existing brain regions can lead to functional divergences [9, 24–26]. These processes can lead to species-specific novel connections not seen in the ancestral circuit or quantitative changes in preexisting projections. However, these evolutionary models of inter-areal cortical connectivity have not been tested quantitatively and in general, mechanisms concerning rapid behavioral divergence remains poorly understood.

The paucity of empirical data to test these models is partially due to technical challenges. Current neuroanatomical methods involve trade-offs between resolution and scalability. Bulk viral tracing of projection patterns can detect novel projections [24, 26, 27] but does not provide quantitative analyses of projection collaterals. At the other extreme, electron-microscopy based connectomes [28–32] provide sub-cellular resolution but remain difficult to scale up for cross-species comparisons in vertebrate brains. Testing models of cortical circuit evolution would require a quantifiable behavioral divergence in a closely-related species, an *apriori* knowledge of a behaviorally-relevant cortical region, and a technique for high-throughput mapping at single cell resolution in many individuals. Therefore, using a striking behavioral novelty in the Alston’s singing mouse (*Scotinomys teguina*) compared to the laboratory mouse (*Mus musculus*), we set out to quantitatively test models of motor cortical evolution using high-throughput barcoded projection mapping of thousands of neurons across multiple animals.

Our focal species is the Alston’s singing mice (*Scotinomys teguina*) – a neotropical, cricetid rodent, emerging as a mammalian model system for studying neural mechanisms of vocal communication [33]. Singing mice produce long, stereotyped, human-audible songs [34–37] that are used for antiphonal (i.e., call-and-response) interactions that exhibit some parallels with human conversational turn-taking [38– 41]. Our previous work, using electrical stimulation, pharmacological inactivation, focal cooling, and chronic silicon probe recordings, has demonstrated that this vocal turn-taking behavior is critically dependent on the orofacial motor cortex (OMC) [37, 42]. Crucially, lab mice (*Mus musculus*), separated from singing mice by around 25 million years [43, 44] (**Fig. 1a**), produce ultrasonic vocalizations (USVs) and there is no clear evidence for turn-taking [41, 45, 46].

**Fig. 1.**
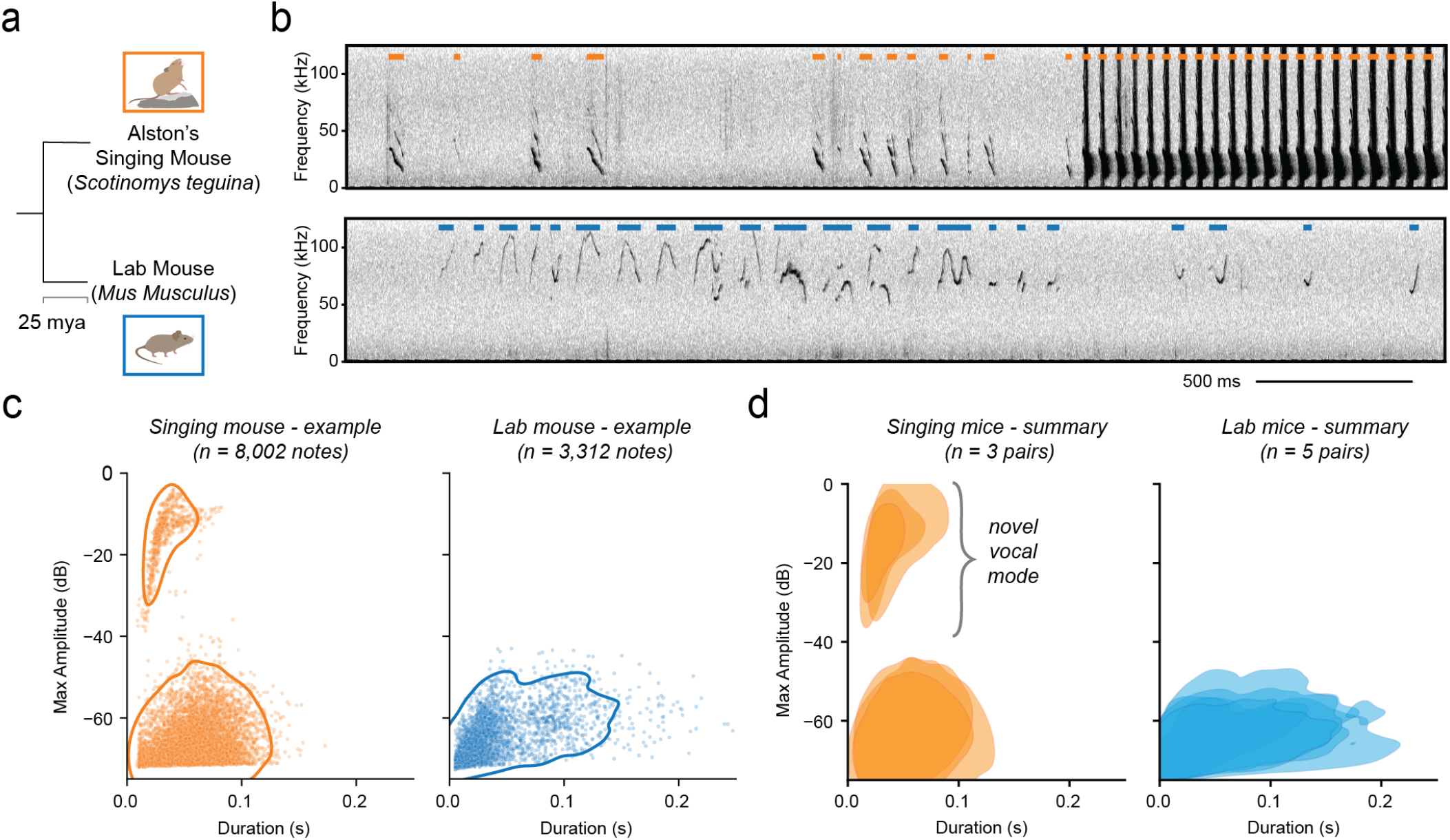
Behavioral novelty in the singing mouse. **(a)** Phylogenetic relationship of the two focal species: singing mouse and lab mouse, which diverged around 25 million years ago (mya) [43, 44]. **(b)** Example spectrogram of singing mice (*top*) and lab mice (*bottom*) vocalizations during male-female social interaction. Orange and blue colored bars mark the duration of each vocalization for the two species. **(c)** Vocalizations plotted in the acoustic space defined by note duration and amplitude from a single pair of singing mice (*left*) and lab mice (*right*). **(d)** Summary of vocalizations from multiple animal pairs (n = 3 pairs singing mice, 5 pairs lab mice). Contours represent areas containing 95% of all notes. Singing mice display two vocalization types (*left*, USV note duration range: 18.5 - 98.5 ms, normalized amplitude range: -71.3 - -54.5 dB., n=11,767 notes; song note duration range: 19.5 – 72.7 ms, normalized amplitude range: -28.3 - -6.8 dB, n=2,046), while lab mice only display a single vocalization type (*right*, USV note duration range: 7.5 – 134.0 ms, normalized amplitude range: -71.7 - -55.5 dB, n=8,385). Ranges represent 5^th^ and 95^th^ percentile values.

In what follows, we first describe that the singing mouse has evolved a novel vocal mode (songs), while retaining ultrasonic vocalizations (USVs) ancestral to rodents. We then utilize the motor cortex-dependent vocal behavior in the singing mouse and lack thereof in the lab mouse [47, 48] to quantitatively test models of motor cortex evolution. Using bulk tracing, serial two-photon tomography, and highthroughput DNA sequencing of thousands of barcoded neurons, we discover selective expansion of OMC projections in the singing mouse to two specific downstream targets: an auditory cortical region (AudR) and the midbrain periaqueductal gray (PAG). Our results suggest that large behavioral divergences over short evolutionary timescales may not require drastic modifications in brain architecture but may instead proceed by modifying statistics of long-range projection patterns.

## Results

We began by quantifying the phenotypic (behavioral) divergence between the two rodent species in identical contexts. We paired adult conspecifics of opposite sex (n = 3 pairs singing mice, 5 pairs lab mice) and allowed them to interact freely in a behavioral chamber for one hour while continuously recording their vocalizations. This paradigm elicited robust vocalizations in both species (n = 13,813 singing mice vocalizations and 8,385 lab mice vocalizations) allowing quantitative assessment of their acoustic features. Consistent with previous reports, we found that lab mice produce USVs in this affiliative behavioral context (**Fig. 1b-d, Supplementary Movie 1**).

In comparison, singing mice vocalizations showed two distinct clusters (**Fig. 1b-d, Supplementary Movie 2**). One cluster consists of loud, stereotyped, song notes while the other cluster corresponds to quieter, more variable vocalizations. The louder notes follow a stereotyped temporal sequence over many seconds (i.e., songs, **Fig. S1**). We previously showed that songs are used for vocal turn-taking, requiring auditory-motor coupling mediated by the orofacial motor cortex [37]. In contrast, the quieter cluster overlaps with the corresponding USV cluster from the lab mice (**Fig. 1c,d**). Given multiple reports of USVs in a variety of rodent species [45, 49] (including murid species such as mouse/rats [50] and gerbils [51]; other cricetid species such as Peromyscus [52], golden hamsters [53]; and others such as guinea pigs [54]), we infer that the USVs represent a vocal mode present in the common ancestor of most rodents. This suggests that the singing mouse has evolved a novel vocal (song) mode that is loud, temporally patterned, and used for vocal interactions [36, 37, 55], while retaining the ancestral USV mode. The stark difference between the two rodent species – specifically motor control for temporal patterning of songs and audio-motor coupling for turn-taking – sets the stage for investigating how orofacial motor cortical circuits might differ in the singing mouse compared to the lab mouse. Building upon previous functional results [37, 42], we wondered whether species-typical differences in vocal behaviors between the lab and singing mouse arise from differential projection patterns of OMC neurons. We tested three alternative models for wiring changes that would result in differential neural circuit function. First, OMC neurons in the singing mouse could project to novel target regions not seen in the lab mouse (**Model 1, Fig. 2a, *left***). Alternatively, OMC neurons could project to the same brain regions in both species, but differ in the strength of innervation within target regions (**Model 2, Fig. 2a, *middle***). Finally, neurons could project to the same brain regions in both species, but differ in the probability that a neuron projects to target brain region (**Model 3, Fig. 2a, *right***).

**Fig. 2.**
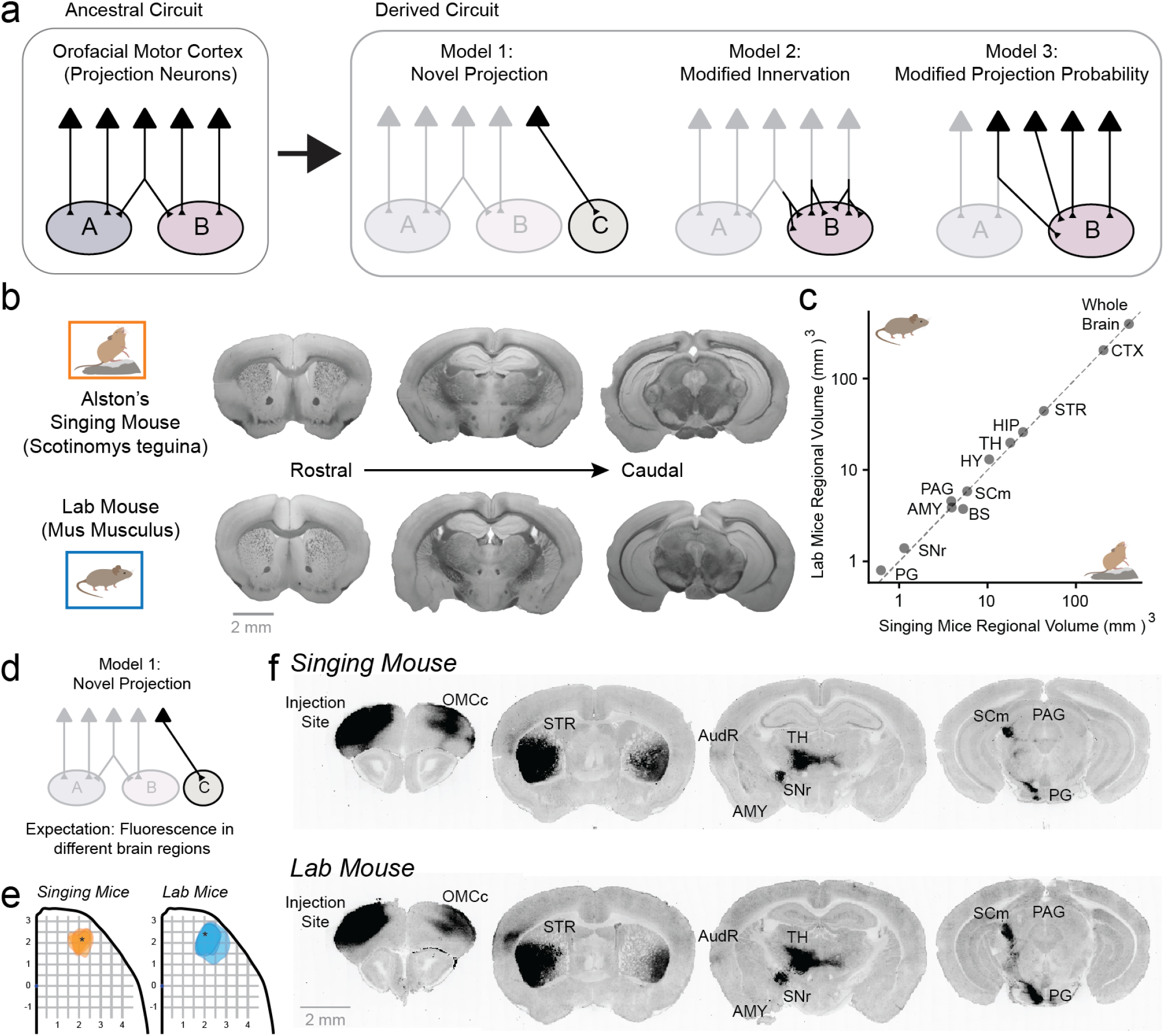
Testing models of motor cortical evolution. **(a)** Three possible models by which orofacial motor cortex (OMC) projections can differ between the two species. **(b)** Comparison of gross anatomical structure of matched brain slices from both species. **(c)** Volume comparisons of selected brain regions in lab mice vs. singing mice. **(d-e)** Testing novel OMC projection targets by viral tracing of axonal projections. Targeting of the homologous OMC regions in both species based on previous studies and overlay of OMC viral spread in singing mice (*left*, n = 3) and lab mice (*right*, n=3). Asterisks mark the center of the injection sites (lab mice: [59, 60]; singing mice: [37]). **(f)** Comparison of brain-wide fluorescence distributions of OMC axonal projections across matched coronal sections from both species. For full dataset, see **Fig S2 and S3**. Abbreviations: cortex (CTX), contralateral orofacial motor cortex (OMCc), striatum (STR), auditory region (AudR), thalamus (TH), substantia nigra (SNr), amygdala (AMY), superior colliculus motor (SCm), periaqueductal gray (PAG), pontine gray (PG).

To test the hypothesis that novel projections exist from OMC in either species, we first measured bulk axonal projection patterns. We injected AAV2.9-CaMKII-tdTomato into the OMC of lab and singing mice (N=3 for each species) and used serial-two photon tomography (STPT,[56]) to obtain an unbiased map of OMC projections throughout the whole brain. Lab and singing mouse brains look nearly identical in gross morphology as illustrated by three representative coronal sections (**Fig. 2b, Supplementary Movie 3 and 4**). We found no significant difference in the overall volume or allometric scaling of OMC or other brain regions among these two species (**Fig. 2c**). Relying on this gross morphological similarity, we aligned all brains to a reference singing mouse brain and compared fluorescence distributions (**Methods, Fig. 2d-f, Fig. S2-S3**). We found that OMC neurons from both species target the same downstream regions (**Fig. 2f, Supplementary Movies 3 and 4**) and match the previously reported motor cortical connectivity in the lab mouse [57, 58]. Thus, we conclude that overall brain architecture and projection patterns of OMC neurons are qualitatively similar between the two species.

In the absence of novel projections, we next wondered whether the anatomical differences between species exist at the single-cell level. While the bulk axonal tracing experiment can map brain-wide projection patterns, the technique cannot resolve quantitative differences in single-neuron projection patterns. Therefore, we employed Multiplexed Analysis of Projections by Sequencing (MAPseq) to map the projection targets of hundreds of motor cortical neurons simultaneously at single-cell resolution [61]. MAPseq experiments involve uniquely tagging hundreds or thousands of neurons with distinct RNA barcodes through injection of a diverse library of barcoded Sindbis virus. The barcodes are expressed and then actively transported into the axonal processes of each labeled neuron. Target regions identified in bulk tracing (**Methods, Fig. 2f, Fig. S4-S5**) are then dissected, and barcodes are extracted and analyzed by high-throughput barcode sequencing (**Fig. 3a**). The abundance of each barcode sequence (i.e., expression level) in each area serves as a measure of the extent of axonal processes of a given cell in a specific target region. MAPseq has been repeatedly validated in multiple studies using multiple methodologies (see **Methods**, [61–67]) but has not been used to test models of neural circuit evolution. MAPseq’s high-throughput and single-cell resolution make it ideally suited to probe quantitative differences in brain-wide projection patterns across species.

**Fig. 3.**
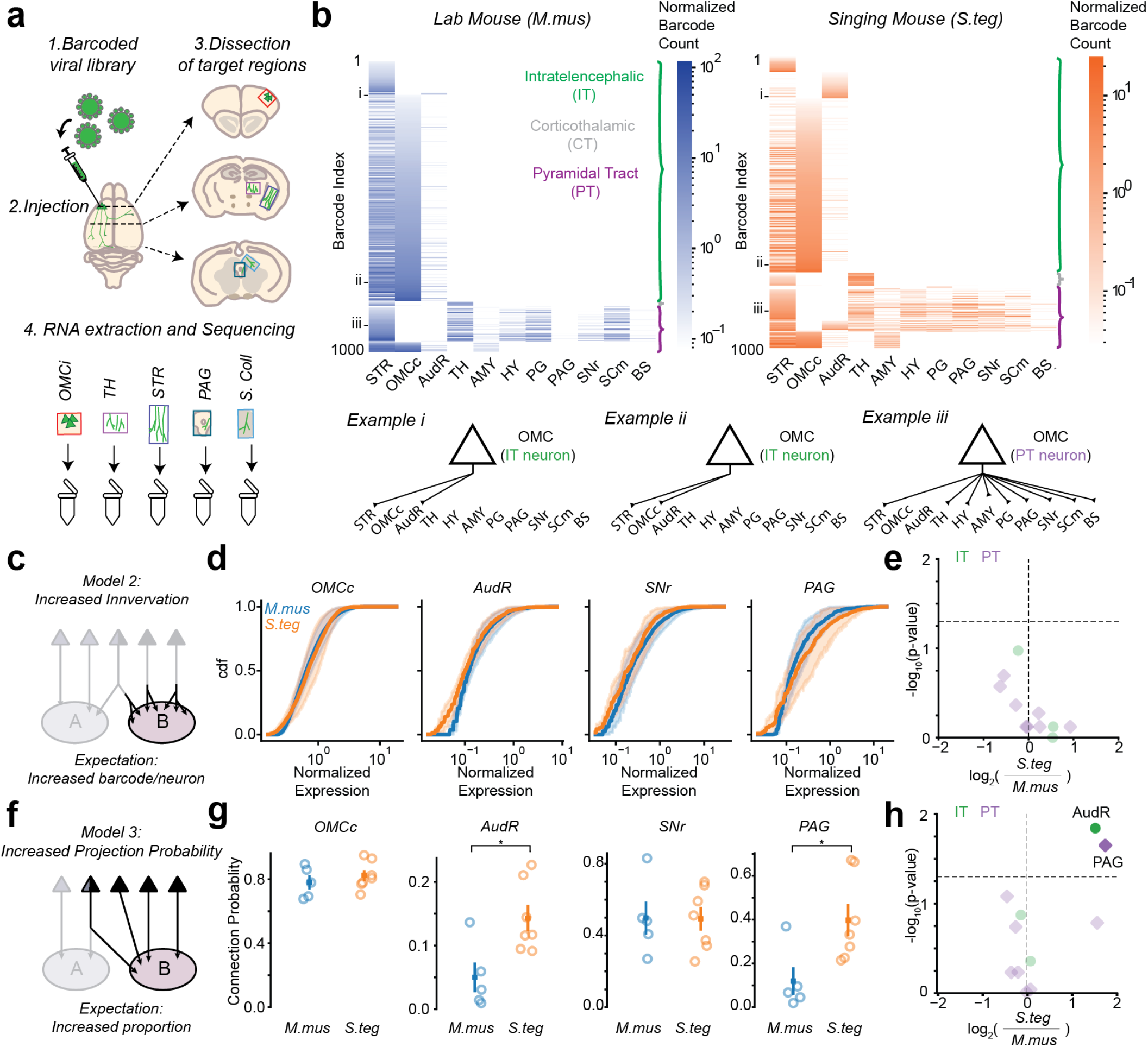
Using MAPseq to test models of projection pattern divergence between lab and singing mice. **(a)** Experimental strategy for high-throughput, single-cell resolution neuroanatomy using molecular barcoding. A Sindbis virus library of unique RNA barcodes are expressed in OMC neurons of each species. Injection sites and target regions are dissected and RNA from each dissected region is extracted and sequenced. **(b)** Single-cell projection patterns of a representative 1000 neurons from an individual lab mouse (*left*) and singing mouse (*right*) of each species showing the major excitatory cell types: IT (green), CT (gray), and PT (purple). Three example (i-iii) single-cell projection patterns of OMC neurons from both species (*bottom*). **(c)** Testing the innervation model by comparing the distribution of barcode expression per neuron within each target region. **(d)** Normalized barcode expression per neuron for individual lab mice (N=5) and singing mice (N=7) are plotted for four selected brain regions. Averages over all individuals with solid lines corresponding to mean and transparent regions corresponding to SEM (OMCc: *M*.*mus*=0.50±0.12, *S*.*teg*=0.73±0.22, p=0.76; AudR, *M*.*mus*=0.13±0.02, *S*.*teg*=0.11±0.02, p=0.11; SNr, *M*.*mus*=0.24±0.06, *S*.*teg*=0.24±0.07, p=0.76; PAG, *M*.*mus*=0.16±0.04, *S*.*teg*=0.31±0.12, p=0.76; Mann-Whitney U test). **(e)** Volcano plot of the negative logarithm of the p-value plotted against the fold change of normalized barcode expression between the two species. No significant differences in the strength of innervation were found within any target region. IT regions are marked by green circles and PT regions are marked by purple diamonds. **(f)** Testing the projection probability model by calculating the proportion of OMC neurons that project to each downstream brain region. Proportions were calculated within the major cell types (e.g. AudR among IT cells, PAG among PT cells). **(g)** Projection probability of OMC neurons to select downstream brain regions. Each data point is an animal and colors indicate species. Solid square and vertical line denotes mean and SEM respectively. (OMCc: *M*.*mus*=0.78±0.04, *S*.*teg*=0.82±0.03, p=0.43; AudR, *M*.*mus*=0.05±0.02, *S*.*teg*=0.14±0.02, p=0.03; SNr, *M*.*mus*=0.50±0.09, *S*.*teg*=0.49±0.07, p=1.00; PAG, *M*.*mus*=0.12±0.06, *S*.*teg*=0.40±0.07, p=0.03; Mann-Whitney U test). **(h)** In the singing mouse, we found a significantly larger proportion of OMC IT neurons projecting to the AudR and OMC PT neurons projected to the PAG in the singing mice compared to lab mice. Horizontal dashed line indicates p = 0.05 (Mann-Whitney U Test)

Using MAPseq, we were able to recover thousands of single-neuron projection patterns in singing and lab mice (**Fig. 3b**, n= 5,114 neurons from 7 singing mice; n=71,704 neurons from 5 lab mice). The expression level of barcodes within infected neurons was similar between species, indicating comparable sensitivity, although the number of recovered unique barcodes per animal differed (normalized barcode expression: *M*.*mus*=0.04±0.02 vs *S*.*teg*=0.13±0.04; unique barcodes per animal: *M*.*mus*=14000±2500, *S*.*teg*=700±100; **Fig. S6**, see **Methods**). Analysis of the MAPseq projection patterns confirmed earlier reports of three major excitatory cortical neurons: intratelencephalic (IT) neurons, which project within the cortex and to the striatum, corticothalamic (CT) neurons, which project from the cortex to the thalamus, and pyramidal tract (PT) neurons, which project from the cortex to the midbrain and brainstem (**Fig. 3b**, [57, 68]).

In the absence of qualitative differences, we tested the hypothesis that the OMC neurons project to the same brain regions in both species, but differ in the strength of innervation within target regions. To determine this, we compared normalized barcode expression per neuron (see **Methods**) across areas and between species (**Fig. 3c-e**). Barcode expression per neuron is proportional to the total axonal volume within the target region much like a fluorescent protein (e.g., GFP) used in conventional anatomical tracing experiments [62, 67]. We did not find any brain region with significant differences in the median barcode expression per neuron (**Fig. 3e**). Therefore, we conclude that there are no significant differences in the strength of innervation of OMC neurons to downstream target regions between the two species.

Finally, we tested the hypothesis that the species difference lies in the probability of OMC neurons projecting to downstream target brain regions. Therefore, we calculated the proportion of OMC neurons that project to specific target regions within each major cell-type (e.g., IT cell for cortical projections; **Fig. 3f-h**). Consistent with previous reports [57, 68], we found that IT neurons from OMC project heavily to the striatum (p(STR) = 0.91±0.02) and contralateral OMC (p(OMCc) = 0.78±0.04), while the PT neurons have strong projections to many regions including the pontine gray (p(PG) = 0.66±0.03), substantia nigra (p(SNr) = 0.50±0.10), and many other subcortical brain regions. For most brain regions (9 out of 11, 82%), we observed no differences in projection probability of IT or PT neurons between the two species. The observed differences were in one IT target (AudR, including primary and secondary auditory cortex) and one PT target (PAG, periaqueductal gray). The projection probability of OMC IT neurons to AudR is significantly higher (180%) in the singing mouse compared to the lab mouse (lab mouse: 0.05±0.05, singing mouse: 0.14± 0.06, p=0.03 Mann-Whitney U test). Similarly, 40% of all OMC PT neurons in the singing mouse projects to the PAG compared to only 12% in the lab mouse, representing a significant (233%) increase in projection probability (lab mouse: 0.12±0.14, singing mouse: 0.40±0.12, p=0.03 MannWhitney U test; **Fig 3h**). We verified that this effect did not depend on the sampling bias by matching the number of neurons from both species (bootstrap down-sampling from the lab mouse dataset, **Fig. S7**). In summary, we find a significant expansion of OMC projections to an auditory cortical region (AudR) and the midbrain PAG in the singing mouse, whose impairments cause specific deficits in vocal communication across a variety of mammalian species [69–74].

So far, we have shown significantly higher projection probabilities of OMC neurons to PAG and AudR in the singing mouse. While this characterizes OMC projections at the population-level, it does not describe the single-neuron projection patterns underlying these differences (**Fig. 4a**). Each neuron has a characteristic projection pattern in a space defined by the number of target regions. For example, an OMC IT neuron in our dataset projecting to potentially three downstream targets can exhibit any one of seven possible motifs (2^3^-1 = 7), constituting its full projectome. The total number of all PT motifs in our dataset is substantially larger (2^8^-1 = 255) making it infeasible to enumerate all the individual motifs. Therefore, we restrict our subsequent analyses to the OMC IT neurons. Specifically, we ask, what is the logic of single neuron level projection patterns underlying the increased projection probability to AudR? In one model, an increase in neural projections to a brain region could arise through increased collaterals, i.e. projections to an additional target region (**Fig 4a, Model 1**). Alternatively, the increase could arise from neurons that project exclusively to that region (**Fig. 4a, Model 2**). In the IT population projecting to the AudR, the distribution of additional collaterals (i.e., the degree distribution) was significantly lower in the singing mice compared to lab mice (**Fig. 4b,c**). This suggests that the expanded OMC projections in the singing mice are preferentially mediated by axons with dedicated projections to AudR.

**Fig. 4.**
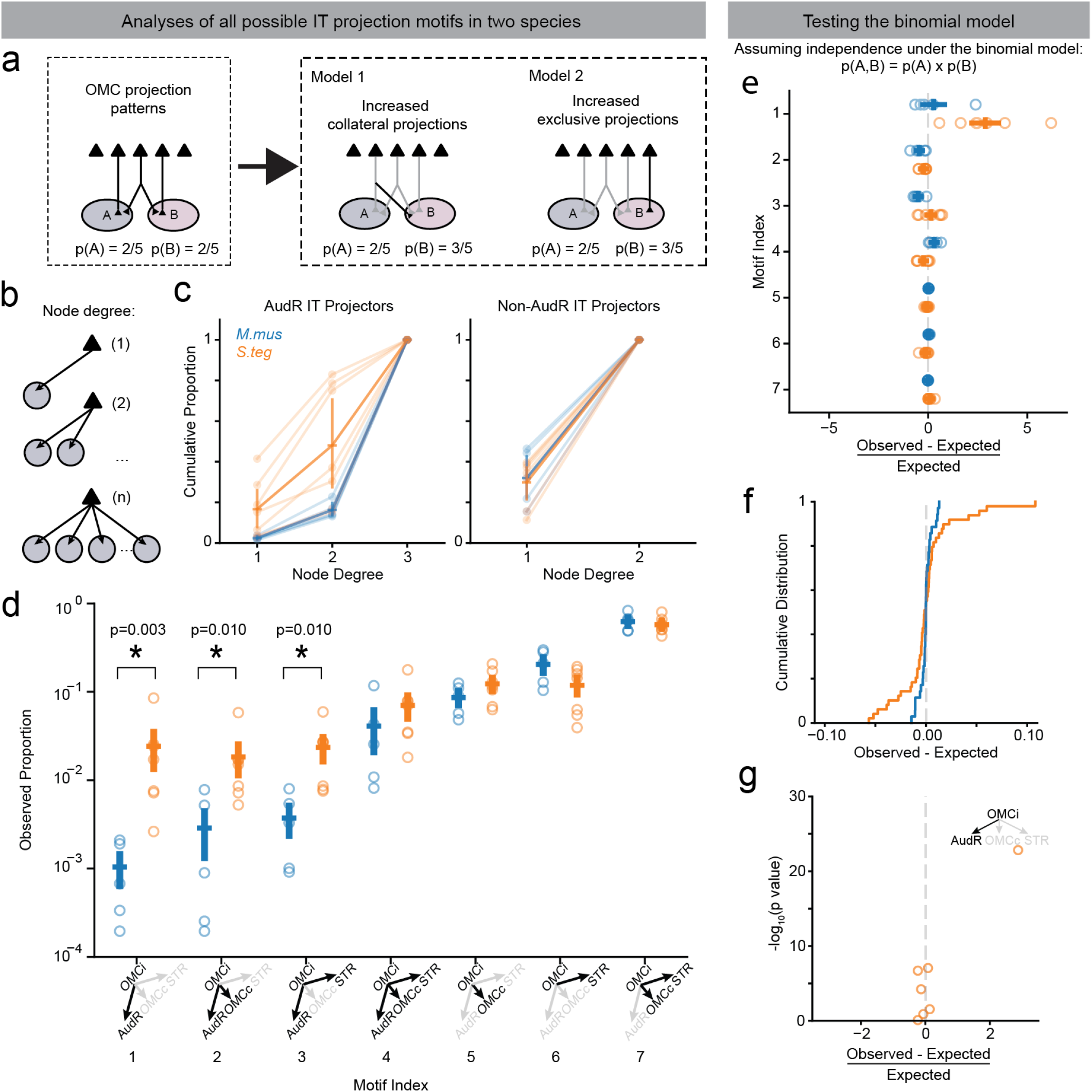
Selective expansion of long-range projection motifs in the singing mice. (**a**) Two models of how increased projection probabilities could evolve from an ancestral state. An increase in projection probability to brain region ‘B’ (e.g., increase from 2/5 to 3/5 in this example) can arise through increased collateral (*center*) or increased exclusive (*right*) projections. Single-neuron resolution projection motif analyses can distinguish between these models of circuit expansion. (**b**) An increased collateral or exclusive projection can be inferred from the node degree distribution of OMC neurons, i.e. the number of targets of an individual neuron. (**c**) Cumulative proportion of node degrees of IT neurons that have AudR as a target (*left*) or do not have AudR as a target (*right*). (**d**) Enumerating the projection probabilities for all seven OMC IT neuron motifs reveals evidence for an expansion of a few select AudR projecting motifs in the singing mice with the largest increase for the exclusive projectors (Motif #1, p value=0.003; Motif #2, p value=0.010; Motif #3, p value=0.010; Mann-Whitney U test). (**e**) Deviation of experimentally measured (observed) motif probabilities from expected values predicted under the binomial model for singing mouse (*orange*) and lab mouse (*blue*). (**f**) Cumulative distribution of differences between observed and expected motif probabilities. All motifs across all individuals for singing mouse *(orange)* and lab mouse *(blue)* are plotted. The singing mouse distribution was significantly different from the lab mouse distribution (p<0.001, F-test). Vertical dashed line denotes the distribution of expected motifs probabilities under the binomial model. (**g**) Binomial test of observed singing mouse motifs probabilities. Each data point corresponds to an individual IT motif.

To verify this possibility, we next enumerated the observed probabilities for all the seven possible IT circuit motifs. While we found increased probability of projections in the singing mouse for most AudR projecting motifs, the single largest expansion was for OMC neurons that exclusively projected to AudR without any axon collaterals in OMCc and STR (p=0.003, Mann-Whitney U test, **Fig. 4d**). By bootstrap resampling, we again confirmed that this result could not be explained by the mismatch in the total number of IT neurons between the two species (**Fig. S7**).

We next wondered whether the multi-areal projection statistics of OMC IT neurons can be explained by a simple binomial model where the probability of projecting simultaneously to multiple regions is simply given by the product of the probabilities of projecting to each region (**Fig. 4e**). Under this assumption of independence, the probability of single OMC neurons projecting simultaneously to two regions (e.g., AudR and OMCc) is expected to be equal to the probability of projecting to AudR multiplied by the probability of projecting to OMCc. For each motif in every animal, we compared the experimentally observed vs. theoretically expected projection probabilities and found overall good agreement with the binomial model predictions (**4e**). Interestingly, the deviation from the binomial model is significantly exacerbated in the singing mice compared to the lab mice (p<0.001, F-Test, **4f**) and the greatest deviance is observed for the motif where OMC neurons project exclusively to AudR without collaterals to OMCc or STR (**4e, g**). We conclude that projection statistics of single OMC neurons can be explained by a simple theoretical framework and a departure from this expectation is a sensitive measure to infer a key locus of evolutionary modification: exclusive OMC projections to AudR in the singing mouse.

Taken together, the reduced degree distribution, the nonuniform expansion of motif probabilities, and the significant deviation from the binomial expectation demonstrate that there are more OMC neurons that project exclusively or almost exclusively to the AudR in the singing mouse. This result suggests that simple uniform increases in neuronal projections across all motifs cannot account for the species-specific differences in single neuronal projection motifs. Rather, a selective expansion of specific long-range motor cortical projection motifs accompanies the evolution of a novel vocal mode in the singing mouse.

## Discussion

In this study, we first demonstrate that songs are a behavioral novelty in the singing mouse compared to the laboratory mouse (**Fig. 1**). We use this striking behavioral divergence between the two rodent species to quantitatively test models of motor cortex evolution (**Fig. 2a**). Using bulk tracing, serial two-photon tomography, and high-throughput DNA sequencing of barcoded OMC neurons at single-cell resolution, we discovered an expansion of motor cortical output to two specific brain regions – auditory cortical region (AudR) and midbrain PAG (**Fig. 2,3**). Using single cell resolution analyses of thousands of cortical neurons, we further show that this expansion is biased toward more exclusive projections to AudR in the singing mouse (**Fig. 4**). Our findings suggest that the selective expansion of existing motor cortical projections may lead to rapid behavioral divergence, providing possible mechanisms for the evolution of enhanced cortical control over vocalizations, an important evolutionary innovation for human language.

These results highlight the utility of the singing mouse model system for testing models of neural circuit evolution. However, the general approach for testing evolutionary models as laid out here: select pairs of related species with large behavioral divergence, perform functional experiments to identify brain regions causally related to the novel behavior, and subsequently investigate neural circuit differences between these species at single-cell resolution, is readily applicable to other model species and clades [75].

Interestingly, the only two brain regions (AudR and PAG) with differential OMC innervations are important for vocal behaviors. Note that we selected downstream OMC targets based on the bulk projection patterns without any obvious behavioral criteria. Thus *a priori* there was no reason for these two brain regions, with well-documented roles in auditory and motor processing during vocalizations, to be detected. The role of the midbrain PAG in vocal production is well documented: PAG is both necessary and sufficient to produce USVs in rodents [71], and lesions in the PAG cause mutism across many species [69, 74]. In addition to motor control circuits, vocal communication also requires the ability to couple vocal response to auditory input from conspecifics, which is dependent on auditory circuits [39, 41, 72]. Closer examination of OMC projections to the AudR (**Fig. 2**) reveals a spatial overrepresentation to the temporal association area (TeA), an understudied region ventral to the primary auditory cortex [72]. This region has been shown to be necessary for socially-motivated maternal behavior [72]. Moreover, OMC and TeA seems to be reciprocally connected (**Fig. S8, Movies S3-S4**). Taken together, we speculate that these bi-directional projections between auditory and motor cortices are crucial for distinct sensorimotor computations involved in vocal turn-taking behavior. Our results do not rule out other loci for circuit modifications. Indeed, understanding how motor cortical control integrates with subcortical [71, 74, 76] and peripheral changes [77] will be necessary to fully account for the behavioral divergence.

Neural circuit differences in adult animals (as reported here) must be set up during development [11, 78]. Our results provide certain constraints on the developmental mechanisms that might generate this species-specific difference in cortical circuit architecture. Despite the apparent molecular complexity in specifying long-range axonal projections, it appears that a simple wiring rule (binomial model with independence) can explain the majority of motif probabilities. The significantly greater probability of the OMC neurons exclusively projecting to AudR in the singing mouse is inconsistent with uniform increase in projection probabilities to all downstream targets. Instead, the selective expansion may occur by direct genetic specification of connectivity strengths in a motif-specific manner. Measuring gene expression along with long-range projection patterns of OMC neurons across ontogeny is necessary to further elucidate the developmental origins of these neural circuit differences.

Large behavioral differences in recently diverged species might be expected to require drastic neural circuit changes. However, recent studies in *Drosophila sp*. show that a small number of quantitative changes – in genomes and in central circuits – may be sufficient to produce a large behavioral divergence in closely-related species [79–81]. Consistent with this emerging view [13**?**] and building on our previous functional studies [37, 42], our results suggest that expansion of vocal repertoire in a mammal may also not require drastic neural circuit changes (**Fig. 2**), but may instead proceed by quantitative modifications of preexisting cortical projection motifs (**Fig. 3,4**). The analytical framework (departure from binomial expectation) described here to identify loci for evolutionary change may even generalize to other model systems. Taken together, quantitative modifications in ancestral circuit connectivity might be a widespread mechanism for allowing behavioral diversification at short timescales.

Perhaps the most striking example of rapid evolution of a behavioral phenotype is the emergence of human language. Language emerged after our divergence from our closest living relatives, chimpanzees, approximately 6 million years ago [82]. Enhanced cortical control over vocalizations has been proposed to be a key feature of human language [39, 83]. We speculate that mechanisms that allow rapid cortex-dependent vocal diversification in rodents might be similar to those required for the evolution of human language in the primate lineage. Genetic mechanisms underlying morphological innovations (e.g., eye structure in animals [84] or prickles in plants[85]) are often conserved over deep evolutionary time – an idea known as deep homology [86]. Similarly, neural mechanisms may also be conserved across a variety of behavioral innovations that require expanded cortical control such as language acquisition. In fact, there is evidence of a strengthened white-matter fiber tract between motor planning and temporal auditory regions in humans as compared to other chimpanzees and macaques [**? ?**]. Elucidating such conserved evolutionary mechanisms is important to understand how neural circuits evolve to create “endless forms most beautiful” [87].

## Acknowledgements

We thank Justus Kebschull, Jessica Tollkuhn, Gregg Castel-lucci, Michael Long, Sanchari Ghosh and members of the Banerjee and Zador laboratories for comments on earlier versions of this manuscript. We also thank Abby Williams, Rhonda Drewes, Thomas Genovese, Sanjeev Kaushalya, and the MAPseq core facility, including Yichen Wu, Diana Ravens, and Corey Freivald, for their technical assistance.

## Funding

NIH BRAIN initiative grant 1RF1NS132046-01 (AB) Searle Scholars Program (AB)

Esther A. & Joseph Klingenstein Fund and the Simons Foundation (AB)

NSF - GRFP (ECI)

## Author contributions

Conceptualization: AB, AMZ

Methodology: ECI, AB, AMZ

Investigation: ECI, CEH, XMZ, HZ, MBD

Resources: HZ, AMZ

Visualization: ECI, AB

Formal Analysis: ECI

Funding acquisition: AB, AMZ

administration: AB, AMZ

Supervision: AB, AMZ

Writing – original draft: ECI, AB

Writing – review and editing: ECI, AB, AMZ

## Competing interests

Authors declare that they have no competing interests.

## Data and materials availability

Requests for resources and reagents should be directed to and will be fulfilled by the Lead Contacts, Arkarup Banerjee and Anthony Zador. This study did not generate new unique reagents. Software for analyzing data is freely available on github: https://github.com/singingmicelab/Iskoetal2024-MAPseq-code. The data analyzed for this study are available upon request from the Lead Contacts.

## Supplementary Materials

**Supplementary Movie 1: Social affiliative vocalizations between male and female lab mice**.

**Supplementary Movie 2: Social affiliative vocalizations between male and female singing mice**.

**Supplementary Movie 3: OMC projection patterns across the whole brain of an aligned lab and singing mouse**.

**Supplementary Movie 4: OMC projection patterns across all aligned replicates**.

**Fig. S1.**
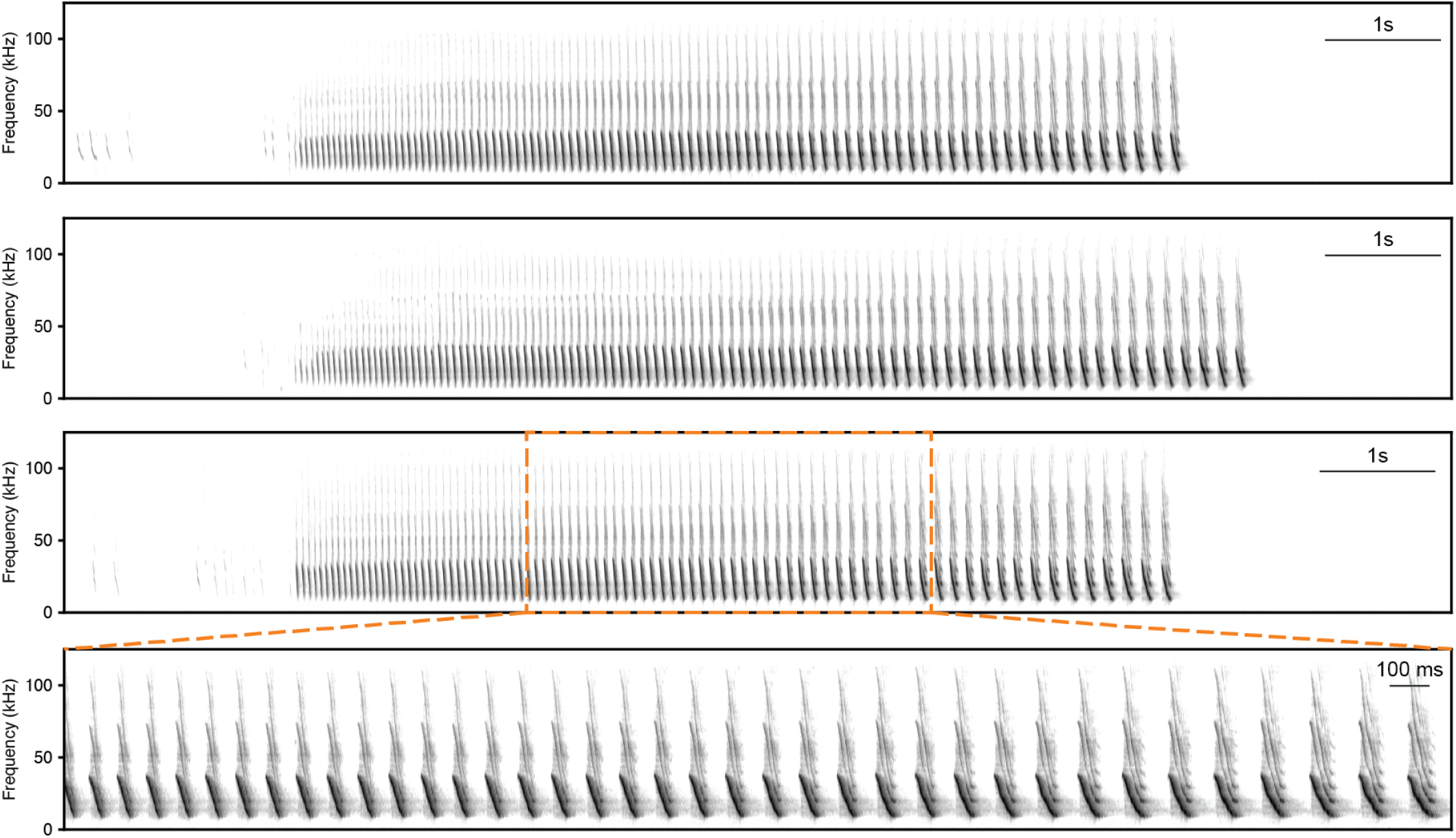
Songs as a behavioral novelty in the singing mice. Three example songs that are composed of a series of frequency-modulated down-sweeps that progress stereotypically over many seconds. Zoomed portion of a song (*bottom*) shows that individual notes are shorter in the beginning and progressively increase as the song progresses.

**Fig. S2.**
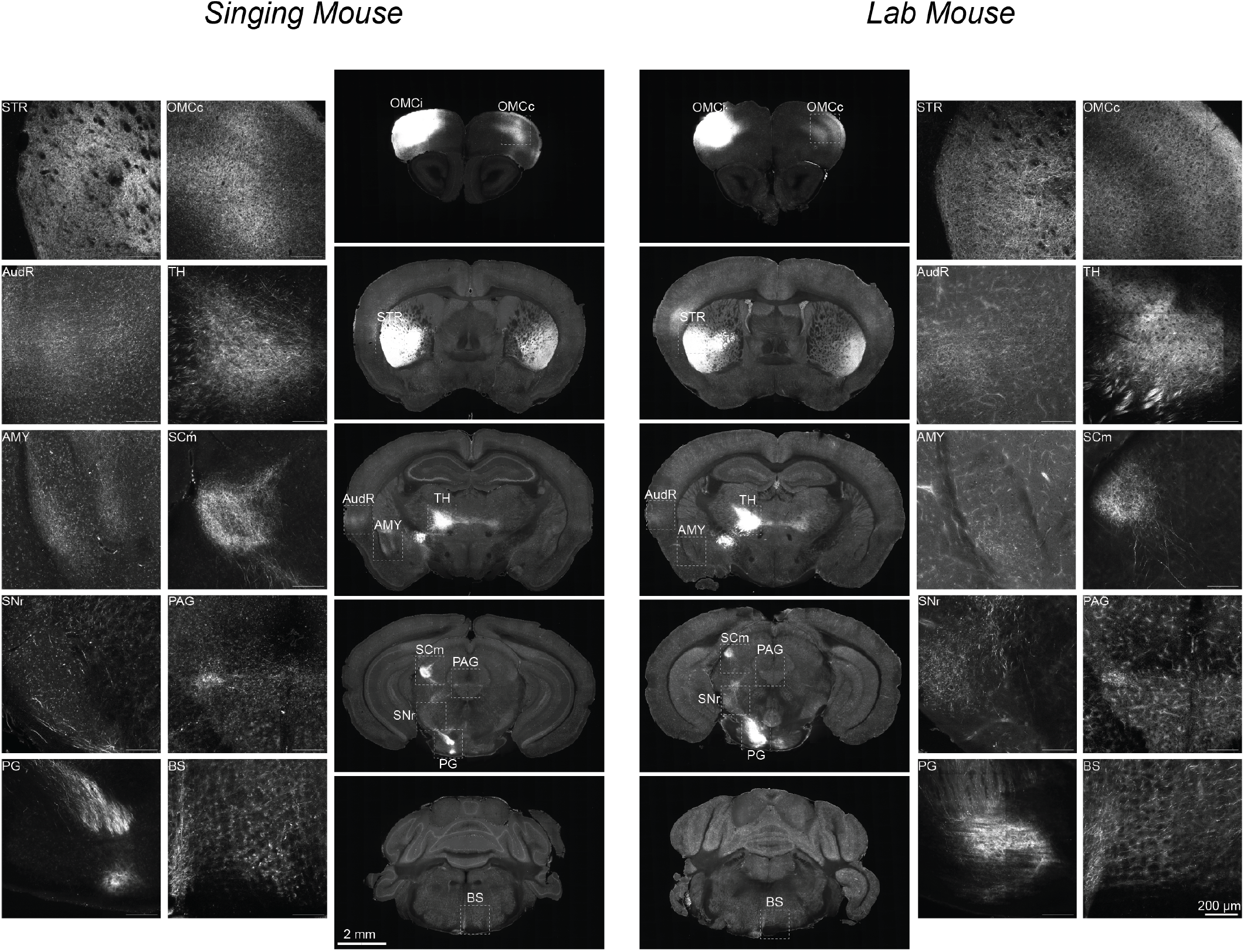
Two-photon microscopy images of OMC neurons projecting to downstream target regions in lab and singing mice. Regions displayed were chosen as MAPseq dissection target regions due to the presence of OMC axonal projections. Abbreviations: Orofacial motor cortex ipsilateral (OMCi), orofacial motor cortex contralateral (OMCc), striatum (STR), thalamus (TH), auditory region (AudR), amygdala (AMY), superior colliculus motor (SCm), substantia nigra (SNr), periaqueductal gray (PAG), pontine gray (PG), brain stem (BS).

**Fig. S3.**
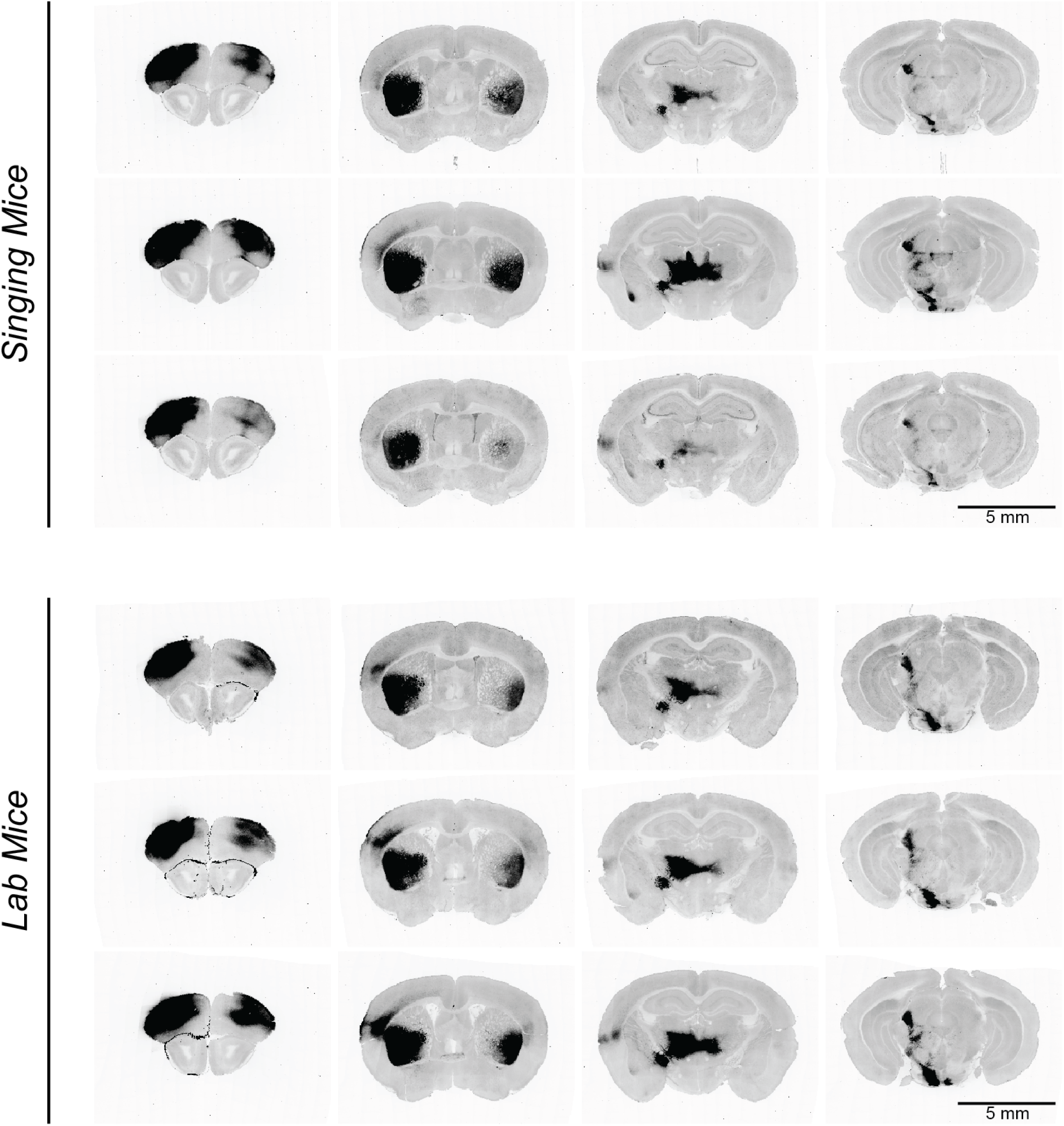
Bulk OMC projections in singing and lab mice across replicates. Representative coronal slices along the rostrocaudal axis (left to right) in individual animals (n= 3) from both species.

**Fig. S4.**
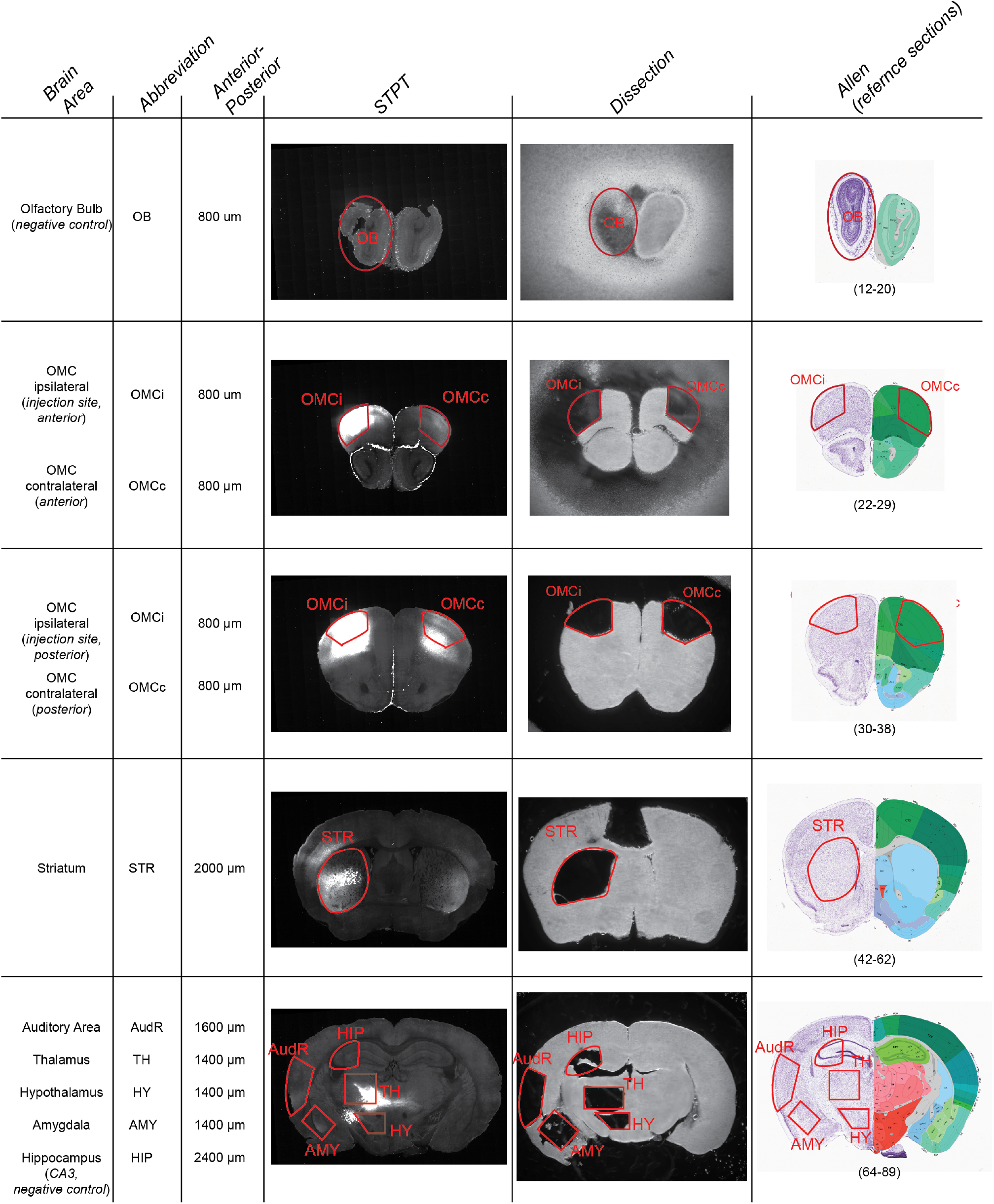
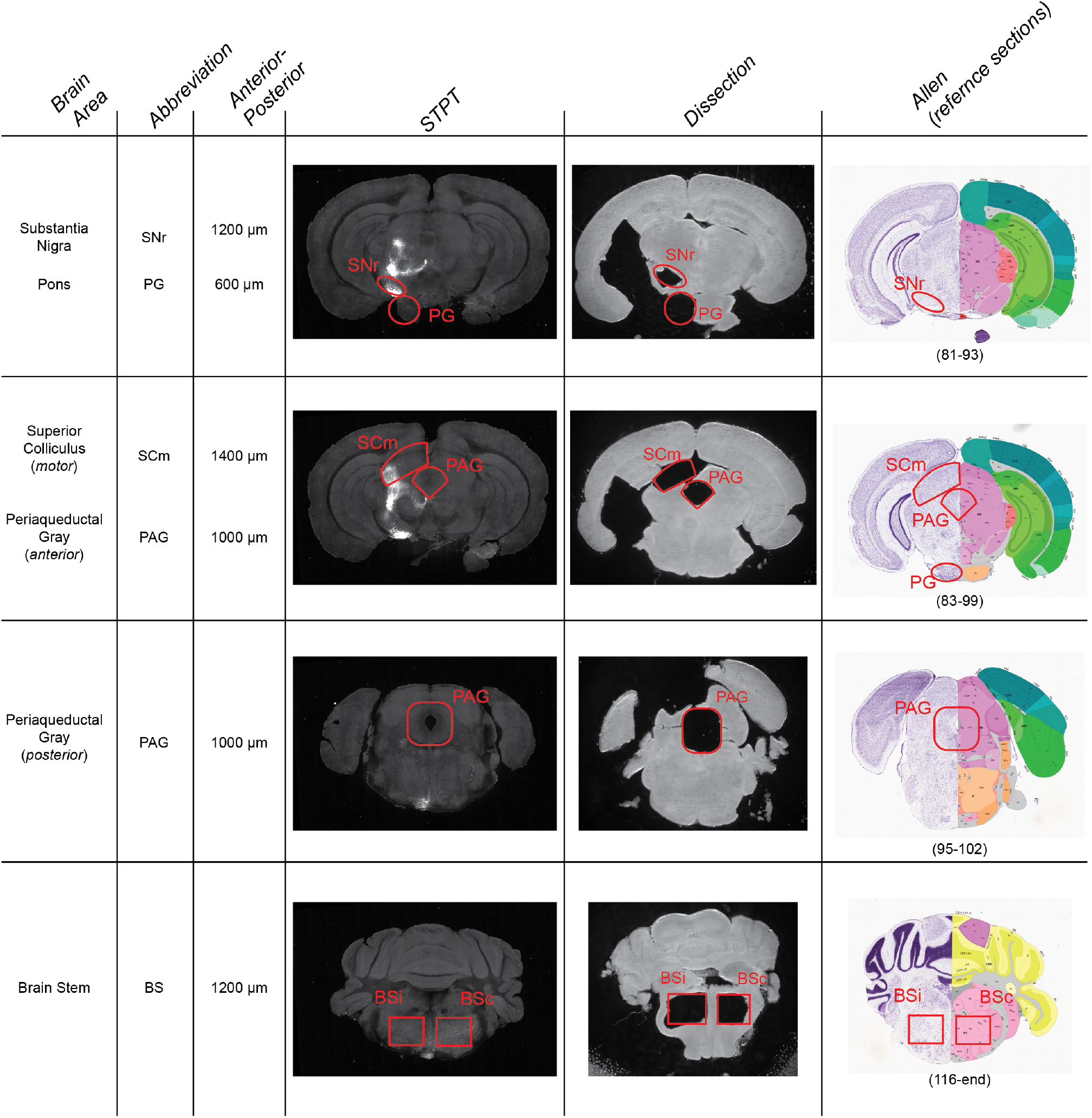
MAPseq dissection guide used for lab mice. Guide used to dissect target regions for MAPseq. STPT data includes representative images from a lab mouse injected with AAV-CaMKII-TdTomatao in OMC (OMCi). Dissection pictures are taken from a single lab mouse MAPseq dissection. Allen references were visually matched to corresponding dissection sections using anatomical landmarks (Allen coronal reference atlas).

**Fig. S5.**
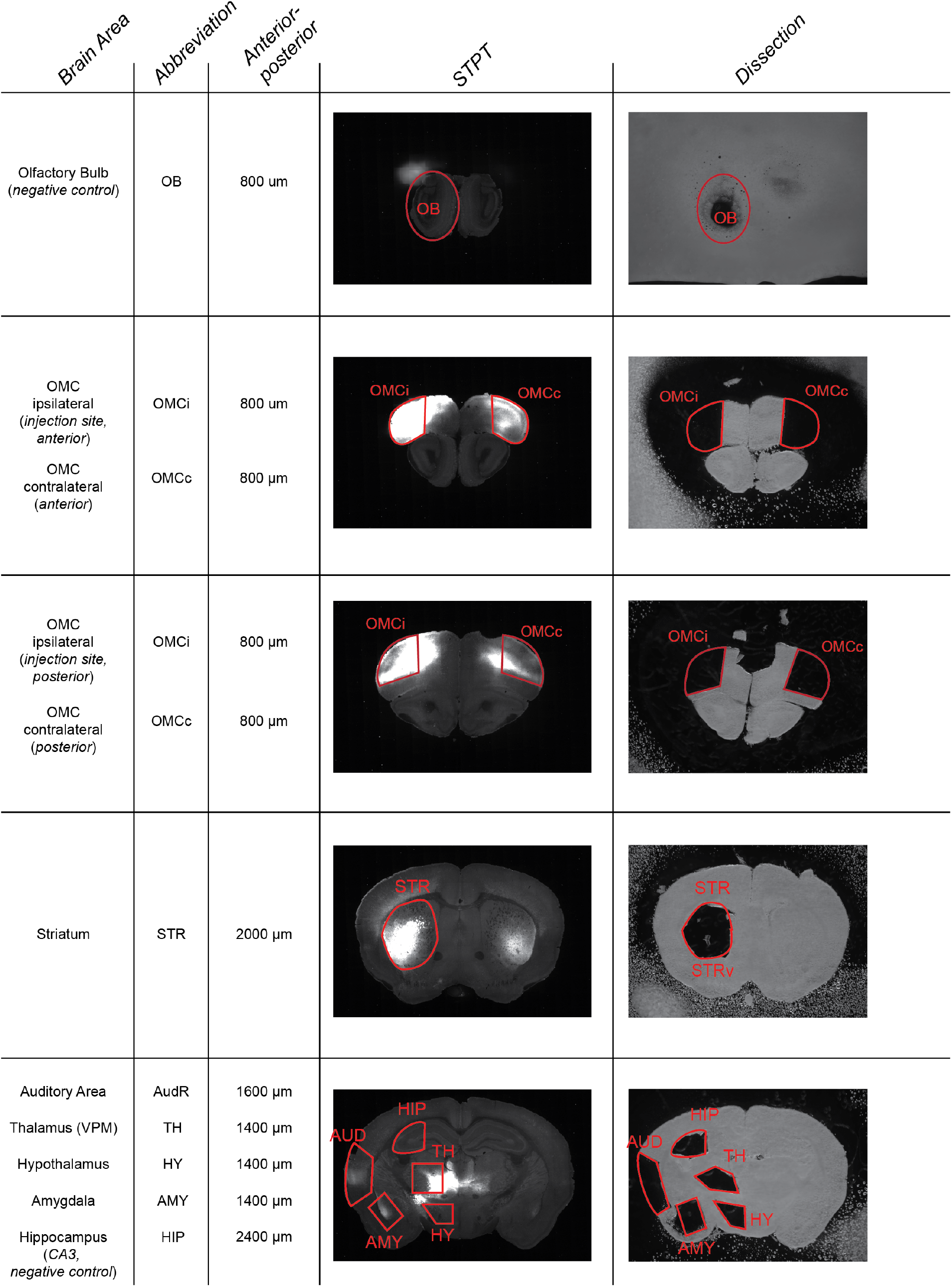
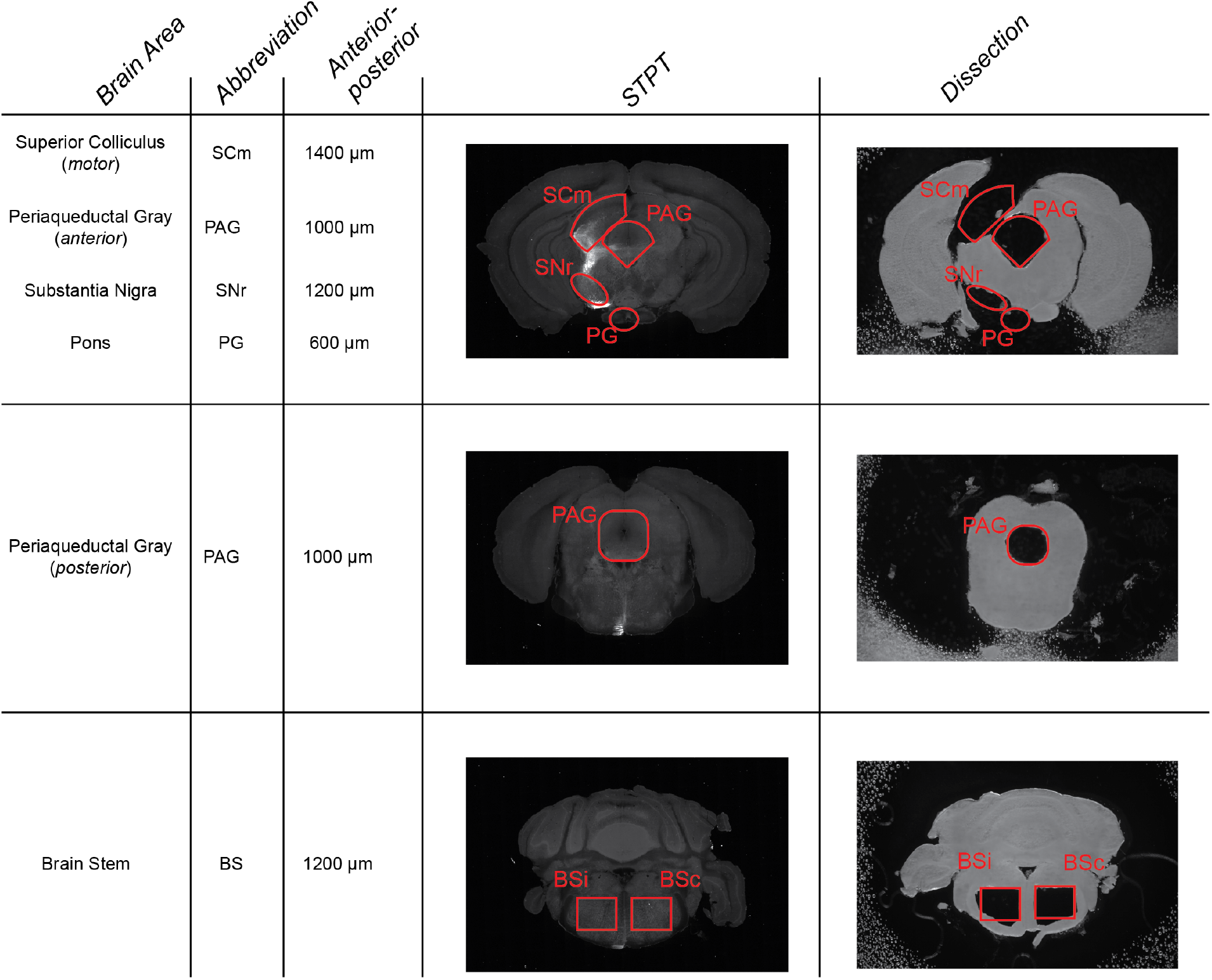
MAPseq dissection guide used for singing mice. Guide used to dissect target regions for MAPseq. STPT data include representative images from a singing mouse injected with AAV-CaMKII-tdTomato in OMC (OMCi). Dissection pictures are taken from a single singing mouse MAPseq dissection.

**Fig. S6.**
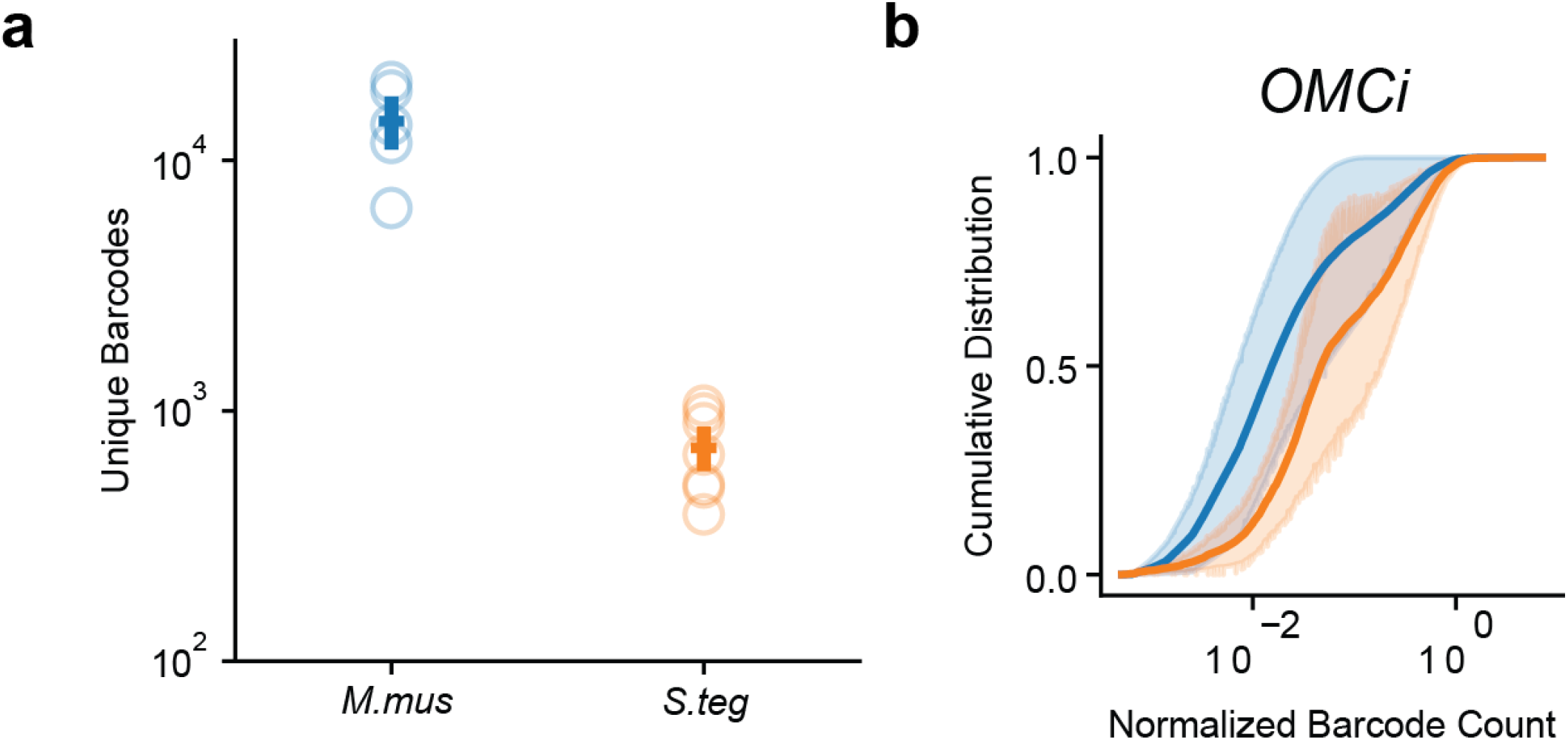
Sindbis infectivity and expression in lab and singing mice. **(a)** Number of unique barcodes recovered in the lab and singing mouse (*M*.*mus*=14000±2500 barcodes per animal, *S*.*teg*=700±100 barcodes per animal). **(b)** Distributions of normalized barcode count in the injection site (OMCi) of lab (blue) and singing mice (orange) (*M*.*mus*=0.04±0.02 vs *S*.*teg*=0.13±0.04).

**Fig. S7.**
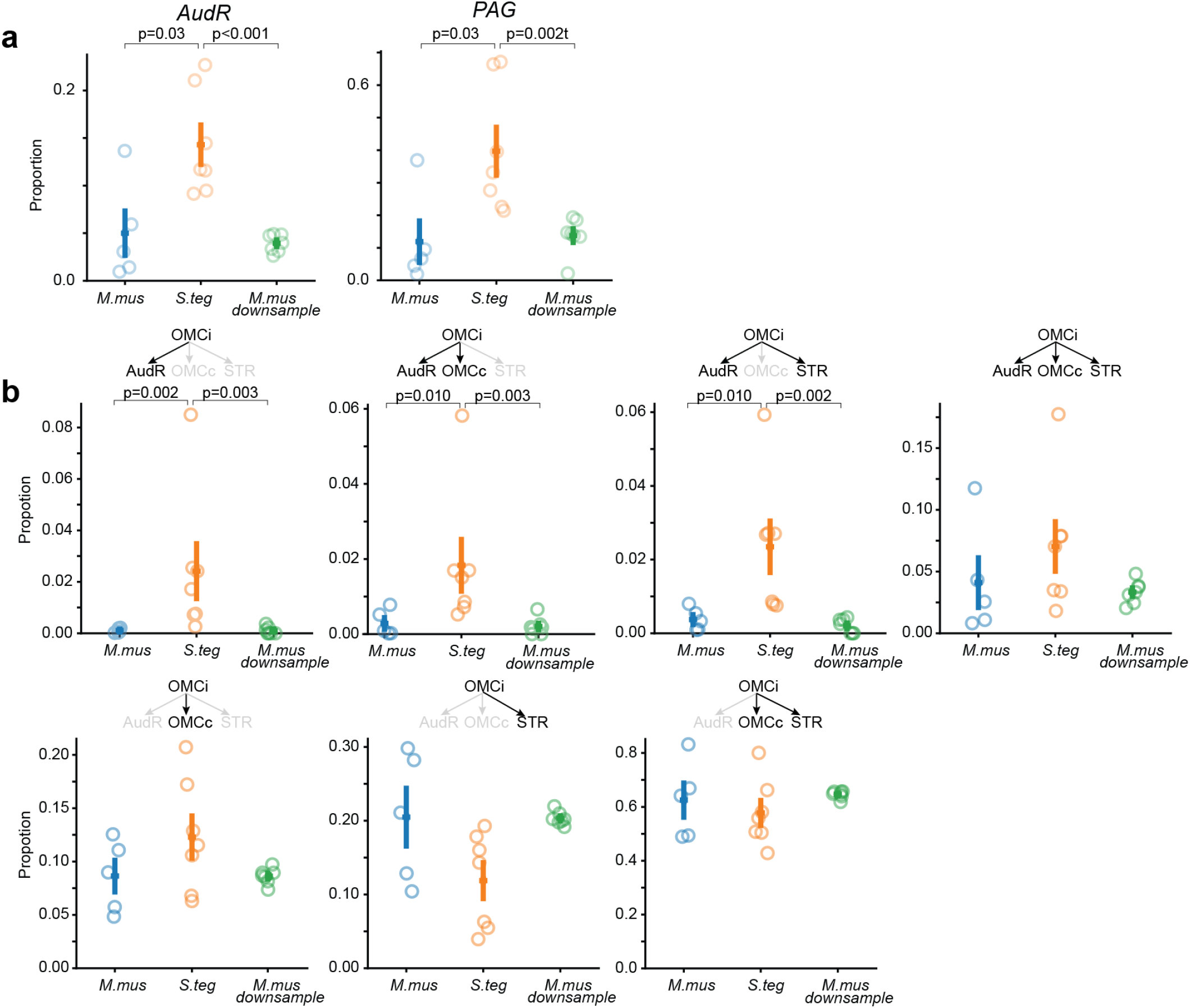
Extended analysis on IT projection and motive proportions. **(a)** Proportion of IT neurons that project to downstream target regions (see **Fig. 3f-h**). Comparison is between lab mice, singing mice, and a downsampled population of lab mice IT neurons. **(b)** Proportion of IT motif in lab mice, singing mice, and a downsampled population of lab mice IT neurons. P values were calculated using a Mann-Whitney U test.

**Fig. S8.**
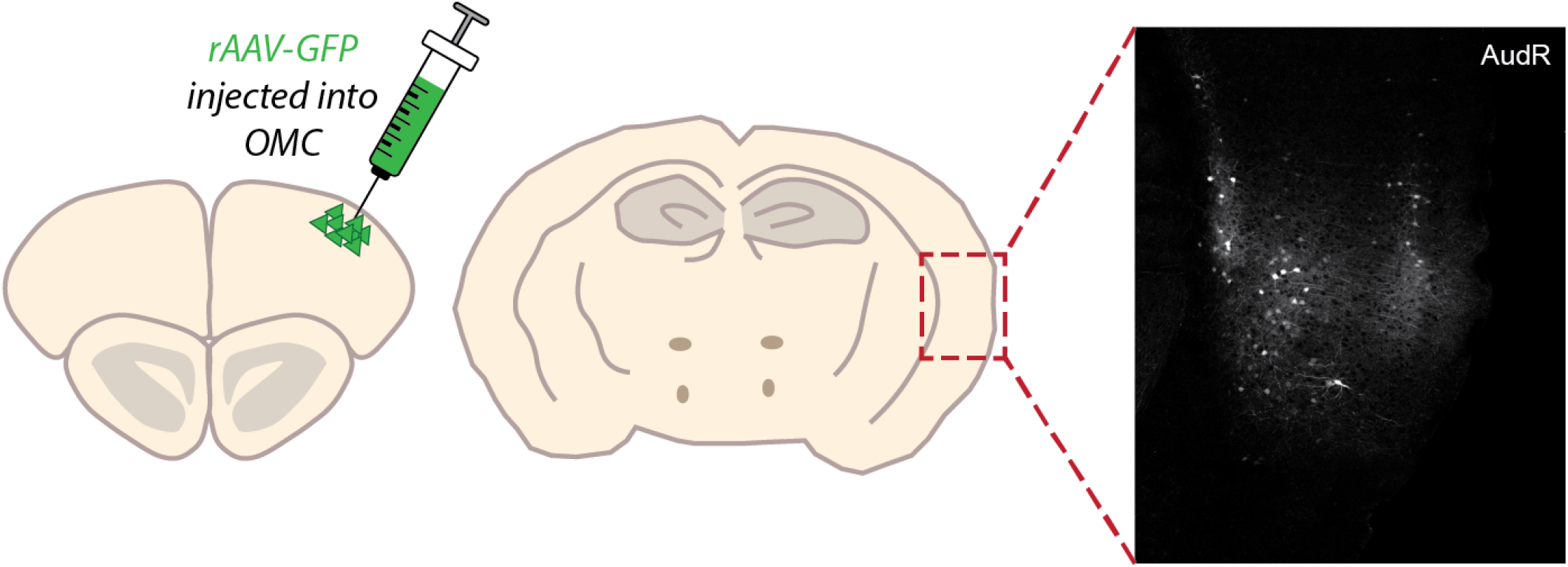
Retrograde viral tracing from OMC reveals synaptic inputs from AudR. Schematic of retro-AAV carrying GFP injected in the singing mouse OMC (*left*). Two photon imaging of the AudR showing neurons projecting to OMC (*right*).

## Methods

### Mouse husbandry

All experiments were approved and conducted in accordance with the Cold Spring Harbor Laboratory Institutional Animal Care and Use Committee. All animals used were adult (>3 months) male and female mice. Lab mice were acquired from Jackson lab (C57BL/6J). Both lab and singing mice colonies were maintained at 20-22°C and a 12:12 L:D cycle.

### Behavioral recordings and Analysis

All mice were singly housed and isolated for at least nine days before social exposure. In advance of recording, the female mouse was placed into a clean cage (Thoren Systems #8, Worcester, MA; 30.80 x 40.60 x 22.23 cm) lined with clean Alpha-pad cotton paper (Shepherd Specialty Papers). After a period of acclimatization (min: 30 mins, max: 12 hours), a non-sibling male of the same species was introduced to the cage with the female. Audio of the pair was recorded for 2 hours using two Avisoft CM16/CMPA microphones positioned above the cage with high and low gain settings and sampling at 250 kHz (digitized with Avisoft UltraSoundGate 116H). Video was recorded using a FLIR Blackfly S high speed USB camera with an Edmund Optics lens at 50 frames per second. The audio and video streams were synchronized using an Arduino Mega and custom code. Biotic sounds were segmented from the audio using USVSEG software for MATLAB (ver. 09r2, [1]) and parameters optimized to the sounds emitted by each species. Additionally, for singing mice, songs and their individual notes were detected using custom code in Python. Segmented sounds were manually curated within a customized spectrogram browser derived from open source MATLAB graphical user interface DeepSqueak [2]. Curation consisted of correcting any biotic sound boundary errors and removal of abiotic false positives. For more accurate amplitude calculations, biotic sounds coinciding with abiotic noise were excluded from analyses. For each curated note, we calculated note duration as the difference between the offset and onset of the max amplitude within a fixed frequency range (10-120 kHz).

### Stereotaxic viral injections

Pre-surgery subcutaneous meloxicam was delivered at a dose of 2 mg/kg. Surgeries were performed on a stereotaxic apparatus under 1-2% isoflurane in oxygen. OMC was localized based on published data (Lab mice: [3, 4]; Singing mice: [5]), which identifies OMC as the area in the motor cortex that when stimulated, results in orofacial muscle contraction (coordinates relative to bregma: [A/P: 2.25 mm, M/L: 2.25 mm]). For mapping of bulk neuronal projections, OMC neurons were targeted to express tdTomato, using a 1:1 mixture of CaMKII-Cre and FLEX-tdTomato viruses (see **Table 1**). Thirty nL of this viral mixture was injected at two depths, 500 and 750 μm ventral to the brain surface, using a Nanoject (Nanoject III, Drummond Scientific, 2 nL/cycle, 15 cycles, at 10 second interval). For MAPseq viral injections, sindbis virus carrying the barcode library [6] was diluted 1:3 in sterile saline. Fifty nL of diluted sindbis virus was injected at 300, 600, and 900 μm ventral to the brain surface at two locations ([A/P: 2.0 and 2.5 mm; M/L: 2.25 mm]) using a Nanoject (Nanoject III, Drummond Scientific, 2 nL/cycle, 25 cycles, at 2 second interval). Mice were transcardially perfused with 4°C PBS and then with 4°C 4% PFA 14 days after injection for STPT or 44 hours after injection for MAPseq experiments.

**Table 1.** Viruses used for bulk fluorescent tracing.

| Virus Name | Titer | Addgene | Serotype |
| --- | --- | --- | --- |
| pENN.AAV.CamKII 0.4.Cre.SV40 | $2.3 \times 10^{13}$ GC/ml | 105558 | AAV9 |
| AAV-FLEX-tdTomato | $2.3 \times 10^{13}$ GC/ml | 28306 | AAV9 |
| rAAV-hsyn-GFP | $2.3 \times 10^{13}$ GC/ml | 50465 | AAV-rg |

### STPT imaging and image processing

STPT imaging was conducted using the protocol established by Ragan et al., 2015 [7] and Kim et al., 2017 [8]. First, brains were embedded in 4% oxidized agarose followed by cross linking in 0.2% sodium borohydrate solution. To image tdTomato projections from OMC in the whole brain, entire coronal planes of the embedded brains were imaged every 50 micros on a TissueCyte 1000 serial two photon tomography microscope (Tissuevision) by tiling and slicing the brain. A chameleon ultra (coherent) 150 fs pulsed laser at 930 nm, Olympus objective (20x, NA=1.0) and 602/70 emission filter was used to image at 1 um/pixel lateral resolution. Image correction and stitching was completed using custom built software [7, 8].

### STPT data analysis

Twenty-fold scaled down images were used for comprehensive and easy visualization between brains. A representative lab and singing mouse brain were chosen as a reference to align brains within species. Additionally, the Allen Brain atlas (2022 CCFv3, [9]) was registered to each brain using the brainreg function with default parameters from the brainglobe suite (https://brainglobe.info/index.html, [10]). Annotated areas from the aligned allen brain atlas were used to determine volumes of major brain regions. To determine the injection site of each animal, a threshold was determined using Otsu’s method. The intersection of the combined annotated primary and secondary motor cortical regions and the thresholded data was used to create a mask of the injection site for each brain. Injection sites of aligned brains were max projected in the dorsal/ventral axis to determine overlap between brains.

### MAPseq tissue processing

After transcardial perfusion, brains were fixed in 4% PFA at 4°C for 24 hours. The brains were then rinsed with PBS and transferred to 300 mM glycine for 24 hours at 4°C. After incubation, brains were embedded in cryo-embedding medium and frozen for sectioning. Brains were cut coronally at 200-250 μm sections for microdissection. Injection sites were identified through brightfield imaging of GFP. During dissection, samples were kept on wet ice during dissection. Target regions were identified through visual landmarks determined from the Allen brain atlas and our STPT data (**Fig. S4-5**). Throughout the procedure, tools and blades were changed between dissection targets and gloves were changed between samples to prevent area and sample cross-contamination. Samples were collected from the olfactory bulb and hippocampus as negative controls. Barcode RNA was extracted, reverse transcribed, and amplified using a published protocol [11, 12]. An illumina sequencing library was generated and sequenced on an Illumina NextSeq500 sequencer. Possible artifacts, including the effect of fibers of passage, viral toxicity, co-infections, double use of a single barcode sequence, and various other sources of false negatives and false positives, have been previously characterized [11, 13]. Notably, although MAPseq, like GFP tracing, does not distinguish fibers of passage, their contribution is minimized by avoiding large fiber bundles during the dissection of target areas.

### MAPseq data analysis

After sequencing, reads were de-multiplexed and the absolute counts of each barcode was determined based on the UMI sequence and error-corrected barcode sequences matched to the sequenced virus library. A matrix of size (number of barcodes) x (number of dissected areas) was constructed where matrix entries corresponded to the absolute counts of individual barcodes in each area. Only barcodes with at least 30 barcode molecules at the injection site and 5 barcode molecules at the maximum target site were used. A barcode was determined to project to a target site if there were at least 4 barcode molecules at that target site. This threshold minimized barcode detection in negative control target regions.

#### Heatmaps and cell type labeling

Normalized barcode counts were calculated from barcode counts normalized to an RNA spike-in of known concentration and quantity in each sample. For heatmaps, 1,000 unique barcodes (i.e. neurons) were randomly sampled from an individual lab or singing mice, and the normalized barcode counts were plotted (**Fig. 3b**) and used to compare Sindbis infectivity between species (**Fig. S6**). Neurons were classified into three major projection, excitatory cell types: IT, CT, and PT. PT cells were first defined as neurons having any projection to a midbrain or brainstem region (HY, PG, PAG, SNr, SCm, or BS). CT cells were defined as having no projections to midbrain or brainstem regions, but having a projection to the thalamus. IT cells were defined as cells having no projections to the midbrain, brainstem, or thalamic regions, and only projections to the striatum and/or cortex.

#### Model 2 - extent of innervation

To eliminate animal-to-animal batch effects, we calculated a normalized barcode expression. For each animal, we took the normalized barcode counts and divided these counts by the mean normalized barcode count at the injecition site (OMCi). Normalized barcode expression was taken as a proxy for the amount of axonal material within each dissected area. The empirical cumulative distribution function of normalized barcode expression for all neurons within a region was plotted to compare species differences. The median value of normalized expression within each region for an individual animal was compared between species, and a Mann-Whitney U test was performed to determine species differences.

#### Model 3 - projection probabilities

To calculate the proportion of neurons projecting to different target regions, the data was binarized (threshold of 4 barcode counts). Proportions were calculated within each major cell type. This accounts for any potential variance in the infectivity across cortical layers, which could bias the relative proportion of major cell types in the dataset. For example, the proportion of OMC neurons projecting to the AudR regions were the number of unique barcodes that had above threshold signal (at least 4 barcode counts) in the AudR region divided by the total number of unique barcodes classified as IT cells. AudR, OMCc, and STR proportions were calculated over the total IT population. HY, PG, PAG, SNr, SCm, and BS proportions were calculated over the PT population. A Mann-Whitney U test was used to calculate significant differences between individuals of each species on a per area basis.

#### Degree and motif analyses

Node degree for a given neuron was defined as the number of projecting areas each neuron has. For instance, if a neuron has projections in areas AudR and STR, this neuron would have a node degree of 2. For degree and motif analyses, only IT cells were analyzed as there were many less PT cells per degree and motif. Motif proportions were calculated within IT cells of an individual mouse. The motif proportions were calculated based on an estimation of the total IT cell population as derived in Han et al., 2018 [13]. In brief, the total IT cell population was adjusted to account for unobserved neurons with projections to uncollected regions or without projections. The total IT cell population can be estimated based on finding the roots of a polynomial derived from observed numbers of neurons to collected regions. The total IT population was calculated for each individual mouse and used as the denominator for determining the observed proportion for each motif. The expected proportion for each motif was calculated by multiplying the probabilities of projecting to individual brain regions according to a binomial model. The difference between the experimentally observed vs. expected proportions for each motif was aggregated for every individual. The extent of deviation from the binomial model was computed for each species and their distributions were compared using an F-test for equality of two variances.

#### Calculating total OMC neuron population

In our MAPseq experiment, the number of IT neurons is defined as the number of neurons that project exclusively to IT regions, i.e. neurons that project to AudR, OMCc, and/or STR. However, the observed number of IT neurons in a MAPseq dataset is a biased sampling of OMC neurons. Due to the nature of the technique, we do not account for neurons that could be part of the OMC IT population but project to regions outside of the regions we sampled. Therefore, the number of OMC IT neurons we count in our dataset is an underestimate of the total OMC IT population. If we want to make reasonable motif predictions under a binomial model, the total OMC IT population count must be adjusted. As detailed in Han et al., 2018 [13], we can derive an estimate for the total OMC IT neurons (N_total_) for each individual animal using our observed counts of how many neurons project to each target region. First, we observe that:

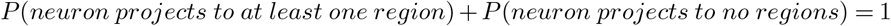

We can also assume that the proportion of neurons that projects to a region is equivalent to the probability that a neuron projects to that region:

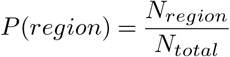

where N_region_ is the number of neurons that project to that region, and N_total_ is the total number of OMC IT neurons. Considering we have 3 IT target regions (OMCc, AudR, and STR), we can combine the above two equations to get the following equation:

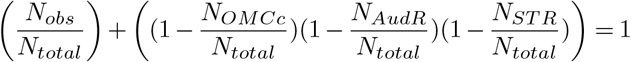

where N_obs_ is the number of unique barcodes (neurons) we recover in our MAPseq experiment, N_OMCc,STR,AudR_ is the number of unique barcodes found to project to an inividual area, and N_total_ is the total number of IT neurons that we are trying to account for. Expanding and rearranging the above equation, we can derive the following quadratic equation:

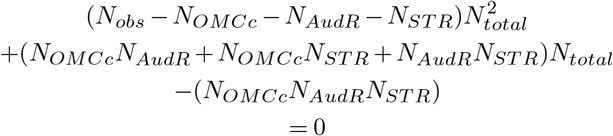

We can then use a roots solver to solve for N_total_. For each animal, we used the observed barcode counts for each region and calculated a quadratic equation using the above formula. We then used a roots solver to find the largest real root. We used this number as the adjusted OMC IT population size (N_total_) and used this N_total_ to calculate the probability that a neuron projects to an individual region. We used these adjusted probabilities to estimate motif proportions expected under the binomial model to give us more accurate predictions.

#### Bootstrap resampling to match the number of neurons across species

Since more unique barcodes were recovered for lab mice, we verified our findings by matching the number of neurons in the singing mice with a downsampled population of neurons in the lab mice. First, all lab mice neurons of the same cell type (IT or PT) were pooled. The pooled neurons were randomly sampled, without replacement, 7 times to exactly match the number of neurons recovered in each of the singing mice individuals. Downstream target proportions and IT motif proportions were calculated for this downsampled population of lab mice neurons. A Mann-Whitney U test was used to calculate IT motif proportion differences between species.

## Notes

### Competing Interest Statement

The authors have declared no competing interest.

### Summary of Updates

Figure 4 panels were updated along with the text in the results section describing figure 4 results. The discussion text was also modified to be more clear.

## Bibliography

1. Pittella, J. E. H. The uniqueness of the human brain: a review. Dement. Neuropsychol. 18, e20230078 (2024). DOI 10.1590/1980-5764-DN-2023-0078.

2. Geschwind, D. H. & Rakic, P. Cortical evolution: judge the brain by its cover. Neuron 80, 633–647 (2013). DOI 10.1016/j.neuron.2013.10.045.

3. Kaas, J. H. The evolution of brains from early mammals to humans. Wiley Interdiscip. Rev. Cogn. Sci. 4, 33–45 (2013). DOI 10.1002/wcs.1206.

4. Mountcastle, V. The evolution of ideas concerning the function of the neocortex. Cereb. Cortex 5, 289–295 (1995). DOI 10.1093/cercor/5.4.289.

5. Krubitzer, L. & Kaas, J. The evolution of the neocortex in mammals: how is phenotypic diversity generated? Curr. Opin. Neurobiol. 15, 444–453 (2005). DOI 10.1016/j.conb.2005.07.003.

6. Krubitzer, L. The magnificent compromise: cortical field evolution in mammals. Neuron 56, 201–208 (2007). DOI 10.1016/j.neuron.2007.10.002.

7. Striedter, G. F. Principles of brain evolution. (2005).

8. Lindhout, F. W., Krienen, F. M., Pollard, K. S. & Lancaster, M. A. A molecular and cellular perspective on human brain evolution and tempo. Nature 630, 596–608 (2024). DOI 10.1038/s41586-024-07521-x.

9. Ebbesson, S. O. The parcellation theory and its relation to interspecific variability in brain organization, evolutionary and ontogenetic development, and neuronal plasticity. Cell Tissue Res. 213, 179–212 (1980). DOI 10.1007/bf00234781.

10. Barker, A. J. Brains and speciation: Control of behavior. Curr. Opin. Neurobiol. 71, 158–163 (2021). DOI 10.1016/j.conb.2021.11.003.

11. Tosches, M. A. Developmental and genetic mechanisms of neural circuit evolution. Dev. Biol. 431, 16–25 (2017). DOI 10.1016/j.ydbio.2017.06.016.

12. Montgomery, S. H., Mundy, N. I. & Barton, R. A. Brain evolution and development: adaptation, allometry and constraint. Proc. Biol. Sci. 283 (2016). DOI 10.1098/rspb.2016.0433.

13. Roberts, R. J. V., Pop, S. & Prieto-Godino, L. L. Evolution of central neural circuits: state of the art and perspectives. Nat. Rev. Neurosci. 23, 725–743 (2022). DOI 10.1038/s41583-022-00644-y.

14. Sakurai, A. & Katz, P. S. Command or obey? homologous neurons differ in hierarchical position for the generation of homologous behaviors. J. Neurosci. 39, 6460–6471 (2019). DOI 10.1523/JNEUROSCI.3229-18.2019.

15. Krubitzer, L., Manger, P., Pettigrew, J. & Calford, M. Organization of somatosensory cortex in monotremes: in search of the prototypical plan. J. Comp. Neurol. 351, 261–306 (1995). DOI 10.1002/cne.903510206.

16. Jorstad, N. L. et al. Comparative transcriptomics reveals human-specific cortical features. Science (2023). DOI 10.1126/science.ade9516.

17. Bakken, T. E. et al. Comparative cellular analysis of motor cortex in human, marmoset and mouse. Nature 598, 111–119 (2021). DOI 10.1038/s41586-021-03465-8

18. Barton, R. A. & Harvey, P. H. Mosaic evolution of brain structure in mammals. Nature 405, 1055–1058 (2000). DOI 10.1038/35016580.

19. Schoenemann, P. T., Sheehan, M. J. & Glotzer, L. D. Prefrontal white matter volume is disproportionately larger in humans than in other primates. Nat. Neurosci. 8, 242–252 (2005). DOI 10.1038/nn1394.

20. Bianchi, S. et al. Dendritic morphology of pyramidal neurons in the chimpanzee neocortex: regional specializations and comparison to humans. Cereb. Cortex 23, 2429–2436 (2013). DOI 10.1093/cercor/bhs239.

21. Benavides-Piccione, R., Ballesteros-Yáñez, I., DeFelipe, J. & Yuste, R. Cortical area and species differences in dendritic spine morphology. J. Neurocytol. 31, 337–346 (2002). DOI 10.1023/a:1024134312173.

22. Krienen, F. M. et al. Innovations present in the primate interneuron repertoire. Nature 586, 262–269 (2020). DOI 10.1038/s41586-020-2781-z.

23. Chartrand, T. et al. Morphoelectric and transcriptomic divergence of the layer 1 interneuron repertoire in human versus mouse neocortex. Science 382, eadf0805 (2023). DOI 10.1126/science.adf0805.

24. Chakraborty, M. & Jarvis, E. D. Brain evolution by brain pathway duplication. Philos. Trans. R. Soc. Lond. B Biol. Sci. 370, 20150056 (2015). DOI 10.1098/rstb.2015.0056.

25. Kebschull, J. M. et al. Cerebellar nuclei evolved by repeatedly duplicating a conserved cell-type set. Science 370 (2020). DOI 10.1126/science.abd5059.

26. Gu, Z. et al. Control of species-dependent corticomotoneuronal connections underlying manual dexterity. Science 357, 400–404 (2017). DOI 10.1126/science.aan3721.

27. Dum, R. P. & Strick, P. L. The origin of corticospinal projections from the premotor areas in the frontal lobe. J. Neurosci. 11, 667–689 (1991). DOI 10.1523/jneurosci.11-03-00667.1991.

28. Winding, M. et al. The connectome of an insect brain. Science 379, eadd9330 (2023). DOI 10.1126/science.add9330.

29. Dorkenwald, S. et al. Neuronal wiring diagram of an adult brain. bioRxivorg 2023.06.27.546656 (2023). DOI 10.1101/2023.06.27.546656.

30. Cook, S. J. et al. Whole-animal connectomes of both caenorhabditis elegans sexes. Nature 571, 63–71 (2019). DOI 10.1038/s41586-019-1352-7.

31. White, J. G., Southgate, E., Thomson, J. N. & Brenner, S. The structure of the nervous system of the nematode caenorhabditis elegans. Philos. Trans. R. Soc. Lond. B Biol. Sci. 314, 1–340 (1986). DOI 10.1098/rstb.1986.0056.

32. Bumbarger, D. J., Riebesell, M., Rödelsperger, C. & Sommer, R. J. System-wide rewiring underlies behavioral differences in predatory and bacterial-feeding nematodes. Cell 152, 109–119 (2013). DOI 10.1016/j.cell.2012.12.013.

33. Banerjee, A., Phelps, S. M. & Long, M. A. Singing mice. Curr. Biol. 29, R190–R191 (2019). DOI 10.1016/j.cub.2018.11.048.

34. Hooper, E. T. & Carleton, M. D. Reproduction, growth and development in two contiguously allopatric rodent species, genus scotinomys. (1976).

35. Miller, J. R. & Engstrom, M. D. Vocal stereotypy and singing behavior in baiomyine mice. J. Mammal. 88, 1447–1465 (2007). DOI 10.1644/06-mamm-a-386r.1.

36. Pasch, B., Bolker, B. M. & Phelps, S. M. Interspecific dominance via vocal interactions mediates altitudinal zonation in neotropical singing mice. Am. Nat. 182, E161–73 (2013). DOI 10.1086/673263.

37. Okobi, D. E., Jr, Banerjee, A., Matheson, A. M. M., Phelps, S. M. & Long, M. A. Motor cortical control of vocal interaction in neotropical singing mice. Science 363, 983–988 (2019). DOI 10.1126/science.aau9480.

38. Pika, S., Wilkinson, R., Kendrick, K. H. & Vernes, S. C. Taking turns: bridging the gap between human and animal communication. Proc. Biol. Sci. 285 (2018). DOI 10.1098/rspb.2018.0598.

39. Castellucci, G. A., Kovach, C. K., Howard, M. A., 3rd, Greenlee, J. D. W. & Long, M. A. A speech planning network for interactive language use. Nature 602, 117–122 (2022). DOI 10.1038/s41586-021-04270-z.

40. Levinson, S. C. Turn-taking in human communication– origins and implications for language processing. Trends Cogn. Sci. 20, 6–14 (2016). DOI 10.1016/j.tics.2015.10.010.

41. Banerjee, A. & Vallentin, D. Convergent behavioral strategies and neural computations during vocal turn-taking across diverse species. Curr. Opin. Neurobiol. 73, 102529 (2022). DOI 10.1016/j.conb.2022.102529.

42. Banerjee, A., Chen, F., Druckmann, S. & Long, M. A. Temporal scaling of motor cortical dynamics reveals hierarchical control of vocal production. Nat. Neurosci. 27, 527–535 (2024). DOI 10.1038/s41593-023-01556-5

43. Steppan, S., Adkins, R. & Anderson, J. Phylogeny and divergence-date estimates of rapid radiations in muroid rodents based on multiple nuclear genes. Syst. Biol. 53, 533–553 (2004). DOI 10.1080/10635150490468701.

44. Steppan, S. J. & Schenk, J. J. Muroid rodent phylogenetics: 900-species tree reveals increasing diversification rates. PLoS One 12, e0183070 (2017). DOI 10.1371/journal.pone.0183070.

45. Sales, G. D. Ultrasound and mating behaviour in rodents with some observations on other behavioural situations. J. Zool. (1987) 168, 149–164 (1972). DOI 10.1111/j.1469-7998.1972.tb01345.x.

46. Holy, T. E. & Guo, Z. Ultrasonic songs of male mice. PLoS Biol. 3, e386 (2005). DOI 10.1371/journal.pbio.0030386.

47. Gan-Or, B. & London, M. Cortical circuits modulate mouse social vocalizations. Sci. Adv. 9, eade6992 (2023). DOI 10.1126/sciadv.ade6992.

48. Hammerschmidt, K., Whelan, G., Eichele, G. & Fischer, J. Mice lacking the cerebral cortex develop normal song: insights into the foundations of vocal learning. Sci. Rep. 5, 8808 (2015). DOI 10.1038/srep08808.

49. Brudzynski, S. M. Handbook of Behavioral Neuroscience (Elsevier, 2018).

50. Portfors, C. V. Types and functions of ultrasonic vocalizations in laboratory rats and mice. J. Am. Assoc. Lab. Anim. Sci. 46, 28–34 (2007).

51. Peterson, R. E. et al. Unsupervised discovery of family specific vocal usage in the mongolian gerbil. bioRxiv 2023.03.11.532197 (2023). DOI 10.1101/2023.03.11.532197.

52. Kalcounis-Rueppell, M. C., Metheny, J. D. & Vonhof, M. J. Production of ultrasonic vocalizations by peromyscus mice in the wild. Front. Zool. 3, 3 (2006). DOI 10.1186/1742-9994-3-3.

53. Floody, O. R. Ultrasonic communication in hamsters. In Handbook of Ultrasonic Vocalization - A Window into the Emotional Brain, vol. 25 of Handbook of behavioral neuroscience, 197–206 (Elsevier, 2018). DOI 10.1016/b978-0-12-809600-0.00019-6.

54. Berryman, J. C. Guinea-pig vocalizations: their structure, causation and function. Z. Tierpsychol. 41, 80–106 (1976). DOI 10.1111/j.1439-0310.1976.tb00471.x.

55. Campbell, P., Pasch, B., Warren, A. L. & Phelps, S. M. Vocal ontogeny in neotropical singing mice (scotinomys). PLoS One 9, e113628 (2014). DOI 10.1371/journal.pone.0113628.

56. Ragan, T. et al. Serial two-photon tomography for automated ex vivo mouse brain imaging. Nat. Methods 9, 255–258 (2012). DOI 10.1038/nmeth.1854.

57. Muñoz-Castañeda, R. et al. Cellular anatomy of the mouse primary motor cortex. Nature 598, 159–166 (2021). DOI 10.1038/s41586-021-03970-w.

58. Vargas, C. D. M. et al. A functional and non-homuncular representation of the larynx in the primary motor cortex of mice, a vocal non-learner. bioRxiv 2024.02.05.579004 (2024). DOI 10.1101/2024.02.05.579004.

59. Komiyama, T. et al. Learning-related fine-scale specificity imaged in motor cortex circuits of behaving mice. Nature 464, 1182–1186 (2010). DOI 10.1038/nature08897.

60. Mercer Lindsay, N. et al. Orofacial movements involve parallel corticobulbar projections from motor cortex to trigeminal premotor nuclei. Neuron 104, 765–780.e3 (2019). DOI 10.1016/j.neuron.2019.08.032.

61. Kebschull, J. M. et al. High-throughput mapping of single-neuron projections by sequencing of barcoded RNA. Neuron 91, 975–987 (2016). DOI 10.1016/j.neuron.2016.07.036.

62. Han, Y. et al. The logic of single-cell projections from visual cortex. Nature 556, 51–56 (2018). DOI 10.1038/nature26159.

63. Huang, L. et al. BRICseq bridges brain-wide interregional connectivity to neural activity and gene expression in single animals. Cell 182, 177–188.e27 (2020). DOI 10.1016/j.cell.2020.05.029.

64. Sun, Y.-C. et al. Integrating barcoded neuroanatomy with spatial transcriptional profiling enables identification of gene correlates of projections. Nat. Neurosci. 24, 873–885 (2021). DOI 10.1038/s41593-021-00842-4.

65. Chen, Y. et al. High-throughput sequencing of single neuron projections reveals spatial organization in the olfactory cortex. Cell 185, 4117–4134.e28 (2022). DOI 10.1016/j.cell.2022.09.038.

66. Zeisler, Z. R. et al. Single basolateral amygdala neurons in macaques exhibit distinct connectional motifs with frontal cortex. Neuron 111, 3307–3320.e5 (2023). DOI 10.1016/j.neuron.2023.09.024.

67. Yuan, L., Chen, X., Zhan, H., Gilbert, H. L. & Zador, A. M. Massive multiplexing of spatially resolved single neuron projections with axonal BARseq. bioRxivorg 2023.02.18.528865 (2023). DOI 10.1101/2023.02.18.528865.

68. Harris, K. D. & Shepherd, G. M. G. The neocortical circuit: themes and variations. Nat. Neurosci. 18, 170–181 (2015). DOI 10.1038/nn.3917.

69. Jürgens, U. The neural control of vocalization in mammals: a review. J. Voice 23, 1–10 (2009). DOI 10.1016/j.jvoice.2007.07.005.

70. Hage, S. R. & Nieder, A. Dual neural network model for the evolution of speech and language. Trends Neurosci. 39, 813–829 (2016). DOI 10.1016/j.tins.2016.10.006.

71. Tschida, K. et al. A specialized neural circuit gates social vocalizations in the mouse. Neuron 103, 459–472.e4 (2019). DOI 10.1016/j.neuron.2019.05.025.

72. Tasaka, G.-I. et al. The temporal association cortex plays a key role in auditory-driven maternal plasticity. Neuron 107, 566–579.e7 (2020). DOI 10.1016/j.neuron.2020.05.004.

73. Marlin, B. J., Mitre, M., D’amour, J. A., Chao, M. V. & Froemke, R. C. Oxytocin enables maternal behaviour by balancing cortical inhibition. Nature 520, 499–504 (2015). DOI 10.1038/nature14402.

74. Schwark, R. W., Fuxjager, M. J. & Schmidt, M. F. Proposing a neural framework for the evolution of elaborate courtship displays. Elife 11 (2022). DOI 10.7554/eLife.74860.

75. Jourjine, N. & Hoekstra, H. E. Expanding evolutionary neuroscience: insights from comparing variation in behavior. Neuron 109, 1084–1099 (2021). DOI 10.1016/j.neuron.2021.02.002.

76. Sweeney, L. B. & Kelley, D. B. Harnessing vocal patterns for social communication. Curr. Opin. Neurobiol. 28, 34–41 (2014). DOI 10.1016/j.conb.2014.06.006.

77. Smith, S. K., Burkhard, T. T. & Phelps, S. M. A comparative characterization of laryngeal anatomy in the singing mouse. J. Anat. 238, 308–320 (2021). DOI 10.1111/joa.13315.

78. Hoke, K. L., Adkins-Regan, E., Bass, A. H., McCune, A. R. & Wolfner, M. F. Co-opting evo-devo concepts for new insights into mechanisms of behavioural diversity. J. Exp. Biol. 222, jeb190058 (2019). DOI 10.1242/jeb.190058.

79. Ding, Y. et al. Neural evolution of context-dependent fly song. Curr. Biol. 29, 1089–1099.e7 (2019). DOI 10.1016/j.cub.2019.02.019.

80. Seeholzer, L. F., Seppo, M., Stern, D. L. & Ruta, V. Evolution of a central neural circuit underlies drosophila mate preferences. Nature 559, 564–569 (2018). DOI 10.1038/s41586-018-0322-9.

81. Auer, T. O. et al. Olfactory receptor and circuit evolution promote host specialization. Nature 579, 402–408 (2020). DOI 10.1038/s41586-020-2073-7.

82. Fitch, W. T. The evolution of speech: a comparative review. Trends Cogn. Sci. 4, 258–267 (2000). DOI 10.1016/s1364-6613(00)01494-7.

83. Simonyan, K. & Horwitz, B. Laryngeal motor cortex and control of speech in humans. Neuroscientist 17, 197–208 (2011). DOI 10.1177/1073858410386727.

84. v. Salvini-Plawen, L. & Mayr, E. On the evolution of photoreceptors and eyes. In Evolutionary Biology, 207–263 (Springer US, Boston, MA, 1977). DOI 10.1007/978-1-4615-6953-4_4.

85. Satterlee, J. W. et al. Convergent evolution of plant prickles by repeated gene co-option over deep time. Science 385, eado1663 (2024). DOI 10.1126/science.ado1663.

86. Shubin, N., Tabin, C. & Carroll, S. Deep homology and the origins of evolutionary novelty. Nature 457, 818–823 (2009). DOI 10.1038/nature07891.

87. Darwin, C. On the Origin of Species (1859).

## Methods Bibliography

1. Tachibana, R. O., Kanno, K., Okabe, S., Kobayasi, K. I. & Okanoya, K. USVSEG: A robust method for segmentation of ultrasonic vocalizations in rodents. PLoS One 15, e0228907 (2020). DOI 10.1371/journal.pone.0228907.

2. Coffey, K. R., Marx, R. E. & Neumaier, J. F. DeepSqueak: a deep learning-based system for detection and analysis of ultrasonic vocalizations. Neuropsychopharmacology 44, 859–868 (2019). DOI 10.1038/s41386-018-0303-6.

3. Mercer Lindsay, N. et al. Orofacial movements involve parallel corticobulbar projections from motor cortex to trigeminal premotor nuclei. Neuron 104, 765–780.e3 (2019). DOI 10.1016/j.neuron.2019.08.032.

4. Komiyama, T. et al. Learning-related fine-scale specificity imaged in motor cortex circuits of behaving mice. Nature 464, 1182–1186 (2010). DOI 10.1038/nature08897.

5. Okobi, D. E., Jr, Banerjee, A., Matheson, A. M. M., Phelps, S. M. & Long, M. A. Motor cortical control of vocal interaction in neotropical singing mice. Science 363, 983–988 (2019). DOI 10.1126/science.aau9480.

6. Yuan, L., Chen, X., Zhan, H., Gilbert, H. L. & Zador, A. M. Massive multiplexing of spatially resolved single neuron projections with axonal BARseq. bioRxivorg 2023.02.18.528865 (2023). DOI 10.1101/2023.02.18.528865.

7. Ragan, T. et al. Serial two-photon tomography for automated ex vivo mouse brain imaging. Nat. Methods 9, 255–258 (2012). DOI 10.1038/nmeth.1854.

8. Kim, Y. et al. Brain-wide maps reveal stereotyped cell-type-based cortical architecture and subcortical sexual dimorphism. Cell 171, 456–469.e22 (2017). DOI 10.1016/j.cell.2017.09.020.

9. Wang, Q. et al. The allen mouse brain common coordinate framework: A 3D reference atlas. Cell 181, 936–953.e20 (2020). DOI 10.1016/j.cell.2020.04.007.

10. Tyson, A. L. et al. Accurate determination of marker location within whole-brain microscopy images. Sci. Rep. 12, 867 (2022). DOI 10.1038/s41598-021-04676-9.

11. Kebschull, J. M. et al. High-throughput mapping of single-neuron projections by sequencing of barcoded RNA. Neuron 91, 975–987 (2016). DOI 10.1016/j.neuron.2016.07.036.

12. Zhan, H., Kebschull, J. & Zador, A. M. MAPseq (multiplexed analysis of projections by sequencing) sample processing protocol. (2021).

13. Han, Y. et al. The logic of single-cell projections from visual cortex. Nature 556, 51–56 (2018). DOI 10.1038/nature26159.

